# Appetitive and aversive cue reactivities differentiate biotypes of alcohol drinkers

**DOI:** 10.1101/2022.07.31.502197

**Authors:** Yu Chen, Chiang-Shan R. Li

## Abstract

Craving reflects the subjective urge to use drugs and can be triggered by both positive and negative emotional states. However, no studies have systematically investigated the relative roles of these mechanisms in the pathophysiology of substance misuse or distinguished the mechanisms in individual vulnerability to substance use disorders. In the current study, we performed meta-analyses of drug cue-elicited reactivity and win and loss processing in the monetary incentive delay task to identify distinct neural correlates of appetitive and aversive responses during cue exposure. We then characterized the appetitive and aversive cue responses in seventy-six alcohol drinkers performing a cue craving task during fMRI. Imaging data were processed according to published routines. The appetitive circuit involved medial cortical regions and the ventral striatum, and the aversive circuit involved the insula, caudate and mid-cingulate cortex. We observed a significant correlation of *β* estimates of cue-elicited activity of the appetitive and aversive circuit. However, individuals varied in appetitive and aversive cue responses. From the regression of appetitive (y) vs. aversive (x) *β*, we identified participants in the top 1/3 each of those with positive (n = 15) and negative (n = 11) residuals as “approach” and “avoidance” biotype, with the others as the “mixed” biotype (n = 50). For clinical characteristics, the avoidance biotype showed higher sensitivity to punishment. In contrast, the approach biotype showed higher levels of sensation seeking and alcohol expectancy for social and physical pressure. The findings highlighted distinct neural underpinnings of appetitive and aversive components of cue-elicited reactivity and substantiated the importance of biotyping substance misuse.

## 1 Introduction

As one of the diagnostic features of alcohol use disorders (AUDs) in DSM-5 (American Psychiatric Association, 2013), craving represents a psychological state in which individuals experience an intense urge to drink. A recent systematic review and meta-analysis suggested that craving plays significant roles in drug use and relapse outcomes and constitutes an important mechanism underlying SUDs (Vafaie and Kober, 2022). Craving often precedes drug use but may also serve to alert individuals to restrain drug-seeking behaviors (Tiffany, 1990). It is of instrumental importance to investigate the psychological and neural processes underlying craving and potential individual differences in these processes (George and Koob, 2022).

Craving can be triggered by alcohol-related stimuli in the environment, memory of the reinforcing effects of alcohol, withdrawal symptoms, and negative mood. Thus, drinkers may experience craving in response to a wide range of environmental and bodily cues – a multidimensional process embodied with different levels of awareness. Thoughts of reinforcing effects goad approach behaviors and engage individuals in alcohol seeking and consumption (King et al., 2014; Le et al., 2022; Zhornitsky et al., 2019). Withdrawal symptoms and emotional distress may also precipitate alcohol use to alleviate physical and mental stress (Fox et al., 2007; Sinha et al., 2009). It is also likely that positive and negative reinforcement both play an instrumental role in motivating alcohol misuse. That is, drinkers may experience a mixed emotional state during craving. Indeed, AUD is known to involve significant individual differences in personality traits, clinical manifestations, and psychiatric comorbidities, and individuals with AUD vary in the psychological, physiological, and neural processes perpetuating alcohol misuse (Clark et al., 2008). A crucial question is whether craving is primarily an appetitive, aversive, or mixed psychological state and whether individuals may experience craving because of these different mechanisms.

Numerous imaging studies have described the neural correlates of cue-induced craving (Hill-Bowen et al., 2021; Jasinska et al., 2014; Sell et al., 1999; Wang et al., 1999; Zeng et al., 2021). A wide swath of brain regions, including the frontoparietal regions, anterior and posterior cingulate, occipital cortex, insula, striatum, amygdala, and thalamus showed higher responses during exposure to drug as compared to neutral cues in individuals with SUDs including AUDs (Kuhn and Gallinat, 2011; Noori et al., 2016; Schacht et al., 2013; Wilson et al., 2018; Zeng et al., 2021; Zhang et al., 2021). Further, the regional, including insular, activities were correlated with subjective urges to use drugs (Bonson et al., 2002; Brody et al., 2002; Colledge et al., 2018; Kilts et al., 2001; Limbrick-Oldfield et al., 2017; McClernon et al., 2005; Myrick et al., 2004; Trotzke et al., 2021; Wang et al., 1999). Substantial research has also revealed neural underpinnings of appetitive and aversive emotional states; for instance, studies of the monetary incentive delay task (MIDT) have characterized the neural correlates of win and loss processing (Bjork et al., 2010; Dhingra et al., 2020; Dhingra et al., 2021). In a recent study, we performed a comprehensive meta-analysis of imaging studies of MIDT (Chen et al., 2022a) and noted significant overlap of the neural correlates of loss processing with those of cue reactivity. The responses of win and loss processing that manifest in the MIDT may provide an opportunity to distinguishing individual appetitive and aversive drug cue reactivity.

To identify the neural correlates shared by cue exposure and valenced motivational states, we thus conducted another meta-analysis focusing cue reactivity on the cue craving task (CCT). From the findings of MIDT and CCT, we characterized the regional activities shared between win and cue exposure and between loss and cue exposure, here referred to appetitive and aversive responses, respectively, in a group of 76 alcohol drinkers. We hypothesized individual variability in appetitive and aversive cue responses and significant differences in clinical characteristics of individuals that showed higher appetitive vs. aversive reactivity and those that showed the opposite.

## 2 Methods

### 2.1 Meta-analyses of CCT

Following the guidelines of “Preferred reporting items for systematic reviews and meta-analyses (PRISMA)”, we searched the literature on PubMed for imaging studies of cue craving task (CCT) with the key words ((alcohol) OR (ethanol) OR (cannabis) OR (THC) OR (joint) OR (cocaine) OR (crack) OR (amphetamine) OR (methamphetamine) OR (nicotine) OR (smoking) OR (smoke) OR (tobacco) OR (cigarettes) OR (heroin) OR (opiates) OR (drug)) AND ((addiction) OR (dependence) OR (abuse) OR (consumption) OR (craving)) AND ((cue) OR (stimulus) OR (stimuli) OR (reactivity)) AND ((fMRI) OR (functional magnetic resonance imaging) OR (neuroimaging)) NOT (review) NOT (meta-analysis). We identified 1,738 studies on February 20, 2022. We also searched on Google Scholar and PsycNet (https://psycnet.apa.org/) using the same key words but found no new studies. A flow-chart for the procedure to arrive at the final sample for meta-analysis of cue-elicited reactivity to drug is shown in **Figure 1**.

**Figure 1.**
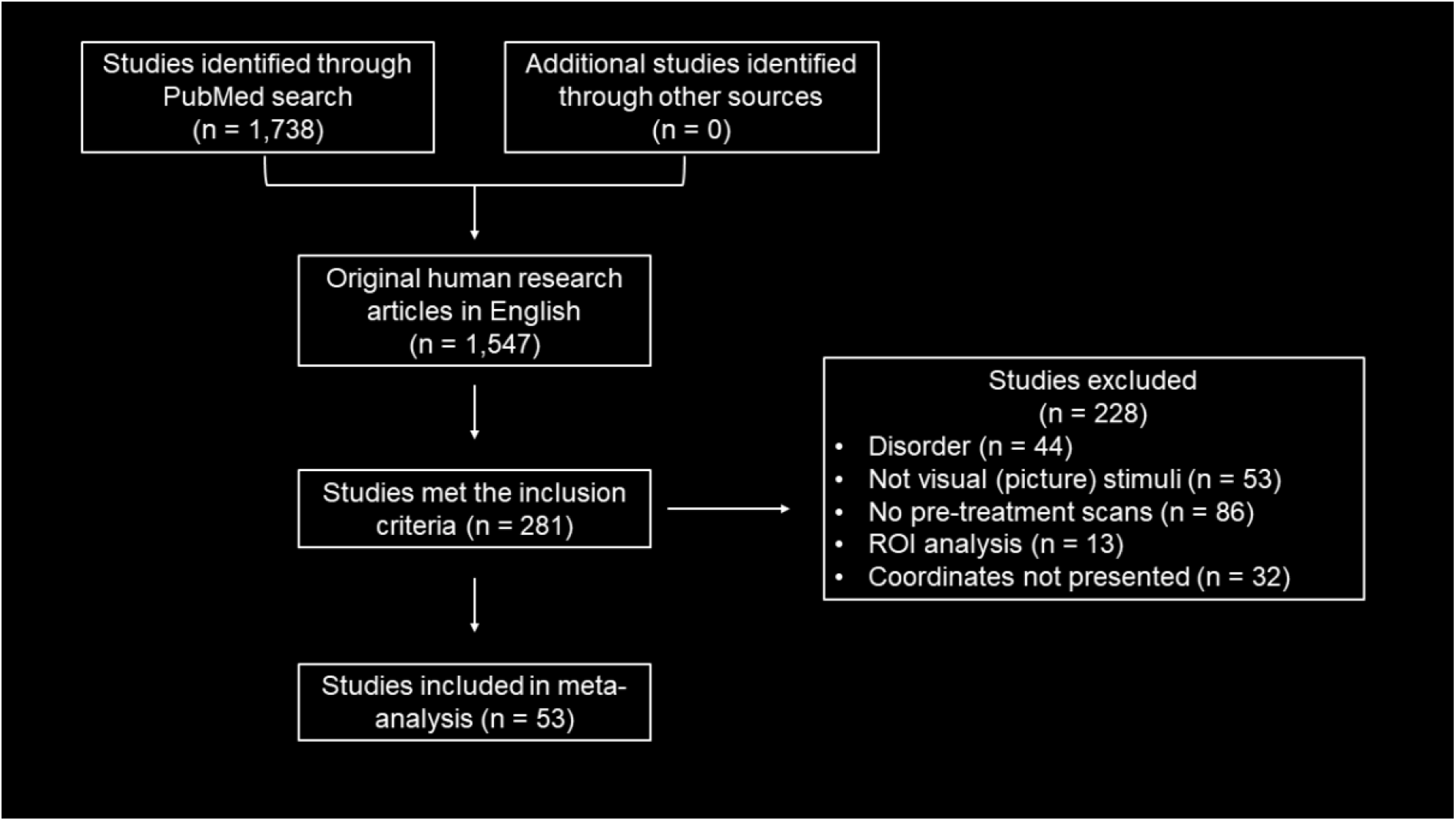
A flow-chart for the procedure to arrive at the final sample for meta-analysis of cue-elicited reactivity to drug, following ‘Preferred reporting items for systematic reviews and meta-analyses (PRISMA).’

Among these 1,738 studies, only peer-reviewed original research articles in English language were included (n = 1,547). Studies of the contrast of drug > neutral cue in smokers, alcohol drinkers, cocaine users, or individuals using other drugs, were included (n = 281). Medication or behavioral treatment studies were included only if the data of pre-treatment scans were available. Studies (n = 44) were removed based on the exclusion criteria, including life-time diagnosis of schizophrenia, depressive disorder, bipolar or manic disorder, psychotic episodes, obsessive-compulsive disorder, or post-traumatic stress disorder; treatment for mental disorders in the past 12 months, use of psychotropic medication; history of or current neurological disorders/brain trauma, or major medical conditions. Studies employing nonvisual stimuli in the CCT (n = 53) or of participants in treatment only (n = 86) were also excluded. Studies (n = 13) performed the regional of interest (ROI) analyses, which violated the assumption of ALE algorithm that each voxel in the entire brain has equal chance of being activated/showing correlation (Muller et al., 2018), were also removed. An additional 32 studies were excluded because the coordinates of the contrast (drug > neutral cue) were not reported. A final pool of 53 studies were included in the current meta-analysis. A complete list of the studies is shown in **Supplementary Table S1**. Notably, in two of these studies, a mixed sample of individuals with different SUDs were included. We converted all foci that were reported in Talairach to MNI space using the Lancaster transformation (Lancaster et al., 2007).

We used the GingerALE software package (version 3.0.2, http://brainmap.org/ale/) to perform ALE meta-analyses on coordinates in MNI space (Eickhoff et al., 2012; Eickhoff et al., 2009; Turkeltaub et al., 2012). The non-additive algorithm was used to reduce the bias of any single experiment (Turkeltaub et al., 2012). The ALE meta-analysis followed four main steps: computation of ALE scores, establishing a null distribution for statistical testing, thresholding, and cluster statistics, as described in detail in the GingerALE Manual (http://brainmap.org/ale/manual.pdf).

We performed the ALE single dataset analysis of drug > neutral cue using a cluster-forming threshold of voxel-level *p* < 0.001, uncorrected. The resulting supra-threshold clusters were compared to a null distribution of cluster sizes established by 1,000 permutations of the data, at an FWE-corrected threshold of *p* < 0.05. We also performed a “Fail-Safe N (FSN)” analysis to evaluate potential publication bias (Acar et al., 2018). We used the R program to generate a list of null studies with no statistically significant activation, all with a number of peaks and a sample size equal to one of the studies in the original meta-analysis. The coordinates of these peaks were randomly drawn from the mask used by the ALE algorithm. We computed the minimum numbers of null studies required in the FSN analysis – 5*k*+10 with *k* denoting the number of studies included in the original meta-analysis (Rosenthal, 1979). Specifically, at least 275 null studies were required. We combined the original and these null studies and repeated the ALE meta-analyses. If the ALE findings remain significant, it means that results are sufficiently robust and are supported by at least the desired minimal number of contributing studies. If adding a minimum of null studies alters the significant results of original ALE analyses, bias due to missing studies (noise) in the meta-analysis is present and the results may not be robust.

### 2.2 Meta-analysis of MIDT

In our previous meta-analysis of MIDT (Chen et al., 2022a), we performed the ALE analyses on single contrasts of anticipation of win vs. neutral (“win anticipation” hereafter), anticipation of loss vs. neutral (“loss anticipation”), win vs. neutral outcome (“win outcome”), or loss vs. neutral outcome (“loss outcome”). Here, we performed ALE analyses with coordinates of contrasts of both win anticipation and outcome (winAO), and both loss anticipation and outcome (lossAO), in combination with a cluster-forming threshold of voxel-level *p* < 0.001, uncorrected. The resulting supra-threshold clusters were compared to a null distribution of cluster sizes established by 1,000 permutations of the data, at an FWE-corrected threshold of *p* < 0.05. We also performed ALE subtraction analyses each to identify regional activities distinct to each contrast (winAO > lossAO and lossAO > winAO). To correct for study sizes (Eickhoff et al., 2011), GingerALE creates simulated data by pooling the foci datasets and randomly dividing them into two groups of the same size as the original data set. An ALE image is created for each new data set and subtracted from the other, with the result compared to the true data. The ALE values were collated across 5,000 permutations to yield an empirical null distribution for statistical inference. A *p*-value was assigned to each voxel based on how many times the difference in the null distribution exceeds the actual difference between the two groups. We applied a threshold of *p* < 0.001 uncorrected with a minimum cluster size of 100 mm^3^ to identify significant differences between any two contrasts. A *Z*-score indicated the size of the differences at each voxel.

### 2.3 Identification of appetitive and aversive cue circuits

To avoid missing any clusters with cue-elicited reactivity in the appetitive and aversive circuits, we used a liberal *p* < 0.05, uncorrected, for the single dataset analyses of drug > neutral, winAO, and lossAO as well as the subtraction analyses of winAO > lossAO and lossAO > winAO. We performed inclusive masking to identify the ROIs.

### 2.4 The empirical study: cue reactivity in drinkers

#### 2.4.1 Subjects, informed consents, and assessments

Seventy-six alcohol drinkers (37 women; age 21-74 years) participated in the study. All participants were required to be physically healthy with no major medical conditions. Those with current use of prescription medications or with a history of head injury or neurological illness were excluded. Other exclusion criteria included current or history of Axis I, including substance (except alcohol and nicotine) use disorders according to the Structured Clinical Interview for DSM-IV (Frances et al., 1995). The study was conducted according to a protocol approved by the Institutional Review Board of Yale University. Written informed consent was obtained from each individual prior to participation.

All participants were evaluated for alcohol use with the Alcohol Use Disorders Identification Test (AUDIT), with a total score ranging from 0 to 40 (Babor et al., 2001). An AUDIT score of 1 to 7, 8 to 14, and 15 or more each suggests low-risk consumption, hazardous or harmful consumption, likelihood of alcohol dependence (moderate-severe alcohol use disorder). Participants were also evaluated for nicotine addiction severity with the Fagerström Test for Nicotine Dependence (FTND) (Heatherton et al., 1991). Ranging from 0 to 10, a higher FTND score indicates more severe nicotine use and dependence. Participants were assessed with the Sensitivity to Punishment and Sensitivity to Reward Questionnaire (SPSRQ), a self-reported instrument that includes 48 yes/no items, comprising a subscale of sensitivity to punishment (SP, 24 items) and a subscale of sensitivity to reward (SR, 24 items) (Torrubia et al., 2001), as well as the UPPS scale to evaluate dimensional impulsivity, including subscales of urgency, lack of premeditation, lack of perseverance, and sensation seeking (Whiteside et al., 2005). In addition, participants were also assessed with the Alcohol Expectancy Questionnaire, an instrument to evaluate alcohol expectancies, including subscales of Positive Global Changes in Experience (GP), Sexual Enhancement (SEXE), Social and Physical Pleasure (SPP), Increased Social Assertiveness (SOCE), Relaxation and Tension Reduction (TRR), and Arousal and Interpersonal Power (PE) (Brown et al., 1987).

#### 2.4.2 Cue-induced alcohol craving task

We employed a cue-induced alcohol craving task (ACT) as described in our previous studies (Le et al., 2020; Wang et al., 2020). Participants viewed alcohol-related or neutral pictures and reported alcohol craving in alternating blocks (**Supplementary Figure S1**). Briefly, a cross was used to engage attention at the beginning of each block. After 2 s, 6 pictures displaying alcohol-related cues (alcohol block) or neutral visual scenes (neutral block) were shown for 6 seconds each. Participants were asked to view the pictures and ponder how they might relate to the images. The pictures were collected from the Internet and independently reviewed by 2 investigators. Alcohol pictures included bar scenes, individuals or a group of people holding or drinking alcoholic beverages, and images of a variety of alcoholic drinks, such as beer, wine, and vodka. Neutral pictures comprised natural sceneries. Participants were asked at the end of each block to report how much they craved for alcohol with rating from 0 (no craving) to 10 (highest craving ever experienced) on a visual analog scale. Each block lasted about 45 s, including time for craving rating. A total of 6 alcohol and 6 neutral blocks took approximately 9 m to complete. Each participant completed 2 runs of the task.

#### 2.4.3 Imaging protocol, data preprocessing and modeling

Briefly, brain images were collected using multiband imaging (multiband factor =3) with a 3-Tesla MR scanner (Siemens Trio, Erlangen, Germany). Data were analyzed with Statistical Parametric Mapping (SPM8, Wellcome Department of Imaging Neuroscience, University College London, U.K.), as in our earlier studies (Wang et al., 2020). Data blocks were distinguished of “alcohol picture” and “neutral picture”, and a statistical analytical block design was constructed for each individual subject using a general linear model (GLM). In the first-level analysis, we constructed for individual subjects a statistical contrast of alcohol vs. neutral blocks.

#### 2.4.4 ROI analyses and group comparisons between biotypes

We computed *β* estimates of cue-elicited activity to alcohol in the ACT within each mask – ROI of the appetitive and aversive cue reactivity circuit identified from meta-analyses – for each participant. We performed a linear regression on the *β* estimates with appetitive and aversive cue reactivity each as the dependent and independent variable. We identified those participants with largest residuals – top 1/3 each of those with positive (n = 15; “approach” biotype) and negative (n = 11; “avoidance” biotype) residuals, with the rest designated as “mixed” biotype (n = 50). We performed a one-way ANOVA to examine whether the biotypes differed in the clinical measures as well as independent-sample *t* tests (two-tailed) to compare the approach and avoidance biotypes, specifically.

## 3 Results

### 3.1 Meta-analyses of cue-elicited reactivity to drug and MIDT and ROI masks

With a cluster-forming threshold of *p* < 0.001 uncorrected and a cluster-level threshold of *p* < 0.05 FWE corrected, cue-elicited reactivity to drug was shown in **Supplementary Figure S2** and the clusters are summarized in **Supplementary Table S2**. A wide array of cortical regions showed higher response to drug vs. neutral cues, including bilateral medial orbitofrontal cortex (mOFC), rostral anterior cingulate cortex (rACC), superior frontal gyri (SFG), precuneus, mid-cingulate cortex (MCC), and posterior cingulate cortex (PCC). The publication bias was evaluated, with 275 additional null studies included in the ALE analyses. The ALE maps evaluated at the same threshold showed a cluster of left mOFC (cluster size: 2,368 mm^3^, MNI coordinates x = −4, y = 50, z = −6, ALE = 0.06; **Supplementary Figure S3**), indicating that the main findings of the original meta-analysis were robust.

The regional activations for winAO and lossAO were shown in **Supplementary Figure S4** and the clusters are summarized in **Supplementary Table S4.** Win anticipation and outcome engaged bilateral parahippocampal gyri, ventral and dorsal striatum, amygdala, anterior insula (AI), thalamus, MCC, PCC, supplementary motor area, precentral gyri, and occipital cortex (OC), and right inferior frontal gyrus (IFG) pars orbitalis. Loss anticipation and outcome engaged similar regional activities (except that the activation of occipital cortex was predominantly in the left hemisphere) and, additionally, the left superior and inferior parietal gyri. These findings are largely in line with previous meta-analyses (Noori et al., 2016; Oldham et al., 2018). The results of ALE subtraction analyses were presented in **Supplementary Figure S5A,** and the clusters are summarized in **Supplementary Table S5**. The contrast “winAO > lossAO” showed higher likelihood of activation in bilateral PCC, whereas the contrast “lossAO > winAO” showed no significant findings. With a liberal threshold of p < 0.05, the subtraction analyses revealed more clusters for both contrasts. The ALE map is shown in **Supplementary Figure S5B,** and the clusters are summarized in **Supplementary Table S5**.

With a p<0.05 to evaluate the findings of both meta-analyses, we overlapped the whole-brain maps of cue-elicited reactivity to drug with “winAO > lossAO” and with “lossAO > winAO”, respectively (**Figure 2A**). Cue-elicited reactivity shared activities with “winAO > lossAO” in bilateral mOFC, rACC, PCC, and OC, as well as right-hemispheric ventral striatum, IFG, and middle frontal gyrus, and left SFG. Cue-elicited reactivity shared activities with “lossAO > winAO” in the left AI and caudate and right MCC. The scatter plot of beta estimates of cue-elicited reactivity is shown in **Figure 2B**, with the three biotypes noted in different colors.

**Figure 2.**
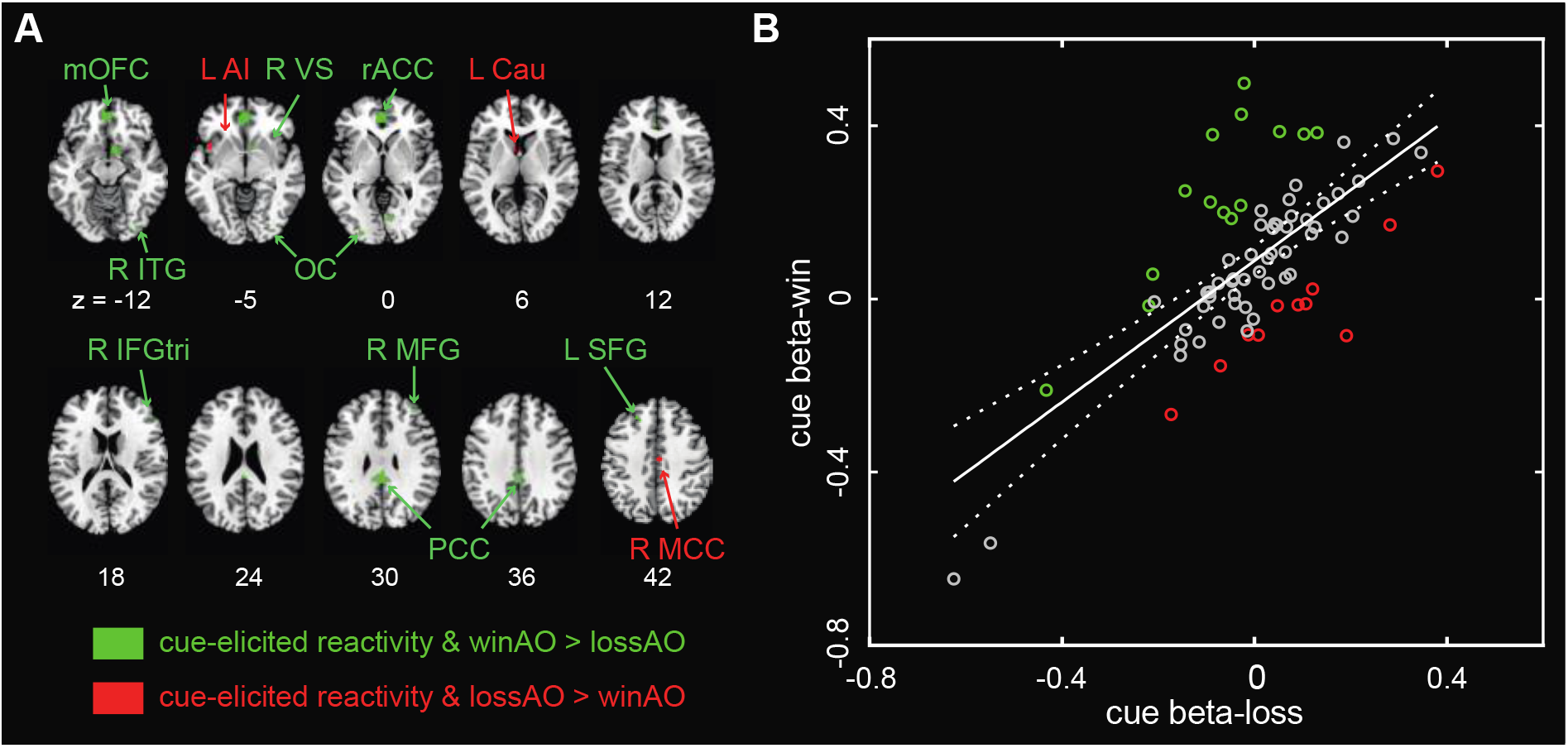
**(A)** Cue-elicited reactivity of the appetitive and aversive circuits. The former involved bilateral medial orbitofrontal cortex (mOFC), rostral anterior cingulate cortex (rACC), posterior cingulate cortex (PCC), and occipital cortex (OC) as well as right-hemispheric ventral striatum (VS), inferior/middle frontal gyrus (IFG/MFG) and left superior frontal gyrus (SFG). The aversive circuit involved the right mid-cingulate cortex (MCC) and left anterior insula (AI) and caudate (Cau). **(B)** Scatter plot of beta estimates of cue-elicited reactivity for approach (green), mixed (grey), and avoidance (red) biotypes. One data point was out of range and not shown.

### 3.2 Differences in clinical characteristics between drinker biotypes

The mean and SD values of demographic and clinical measures of drinker biotypes are presented in **Table 1**. Independent-samples *t* tests showed significant group differences (approach vs. avoidance) in the SPSRQ subscore of punishment sensitivity (7.40 ± 4.39 vs. 13.00 ± 5.78; *t* = −2.81, *p* = 0.010), UPPS subscore of sensation seeking (36.87 ± 5.94 vs. 30.60 ± 7.14, *t* = 2.39; *p* = 0.026), and AEQ subscore of social and physical pressure (24.00 ± 3.49 vs. 19.82 ± 6.66; *t* = 2.08, *p* = 0.048). No significant group differences were found in other demographic or clinical measures (*p’s* ≥ 0.204).

**Table 1.**
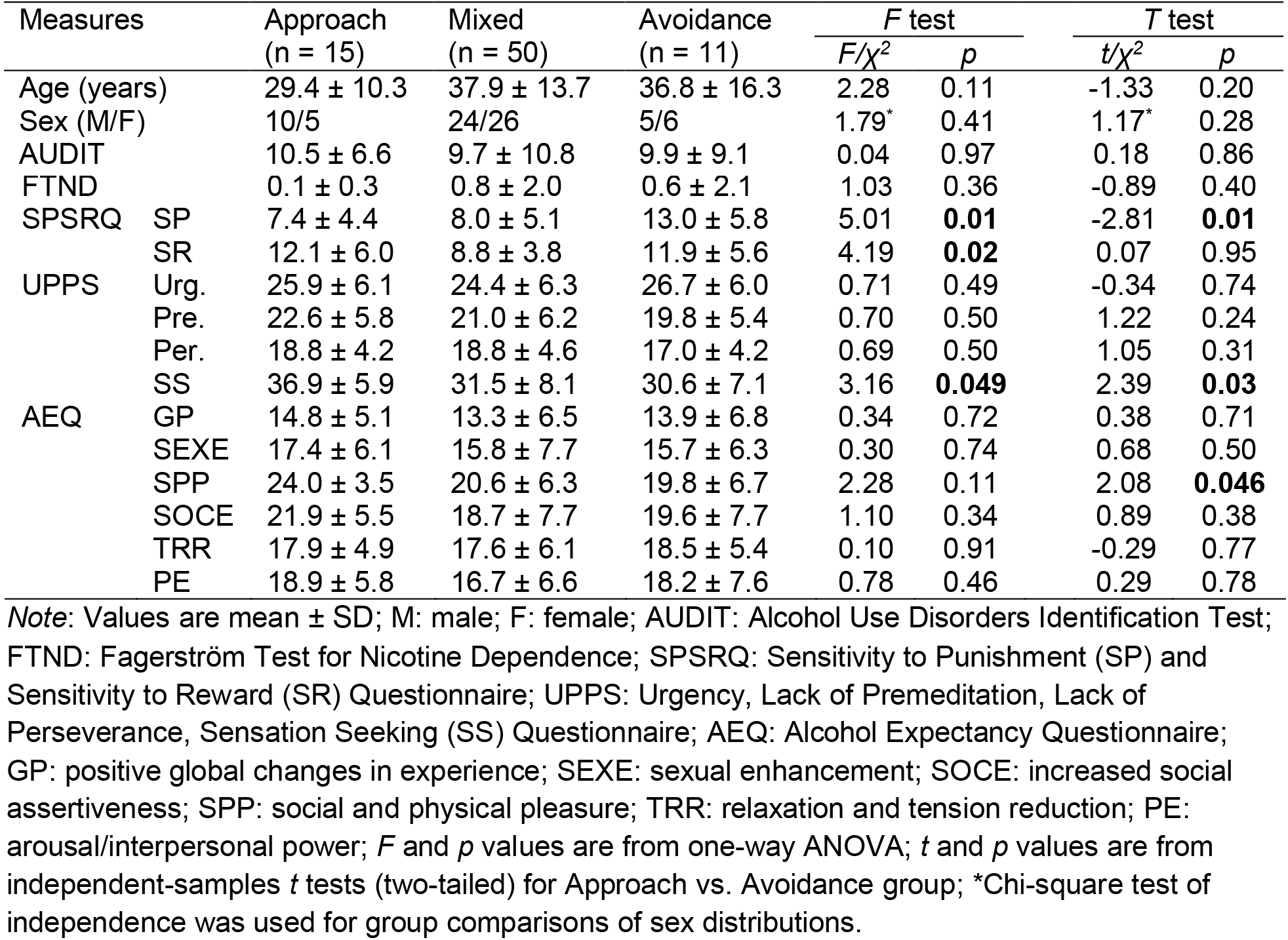
Demographic and clinical measures of drinker biotypes

We also performed one-way ANOVA to examine the differences in these measures with an additional biotype included – mixed biotype. The biotype effect was significant for SPSRQ punishment (*F* = 5.01, *p* = 0.009) and reward (*F* = 4.19, *p* = 0.019) sensitivity and UPPS sensation seeking (*F* = 3.16, *p* = 0.049), but not for AEQ social and physical pleasure (*F* = 2.28, *p* = 0.109). We further performed the post-hoc LSD tests. The avoidance biotype (13.00 ± 5.78) showed significantly higher SPSRQ punishment sensitivity than approach (7.40 ± 4.39; *p* = 0.007) and mixed (7.96 ± 5.06; *p* = 0.004) biotypes, while the latter two showed no significant differences (*p* = 0.707). The mixed biotype (8.80 ± 3.81) showed significantly lower SPSRQ reward sensitivity than approach (12.07 ± 5.97; *p* = 0.018) and avoidance (11.91 ± 5.63; *p* = 0.045) biotypes, while the latter two showed no significant differences (*p* = 0.931). The approach biotype (36.87 ± 5.94) showed significantly higher levels of sensation seeking than mixed (31.54 ± 8.13; *p* = 0.021) and avoidance (30.60 ± 7.14; *p* = 0.048) biotypes, while the latter two showed no significant differences (*p* = 0.723).

## 4 Discussion

To our knowledge, this is the first study to characterize individual differences in the physiological and neural mechanisms with respect to appetitive, aversive, or mixed nature of the psychological state during cue-evoked craving. The medial prefrontal cortex and VS showed appetitive responses and the AI, caudate and MCC showed aversive responses to alcohol cues, consistent with earlier findings (Galandra et al., 2018; Li et al., 2022; Pessiglione and Delgado, 2015; Zhornitsky et al., 2021). By distinguishing appetitive and aversive components of cue reactivity, we identified biotypes of alcohol drinkers, with the approach and avoidance biotype each showing higher appetitive and aversive activity, respectively, relative to the regression mean. The approach relative to avoidance biotype showed lower sensitivity to punishment but higher levels of sensation seeking and expectancy for drinking to increase social and physical pleasure. A mixed biotype, comprising most drinkers in the sample, demonstrated a mixed motivational state during cue exposure and intermediate levels of these clinical traits. We discussed the main findings below.

### 4.1 Shared neural correlates of cue exposure and valenced motivational states

The appetitive circuit involved bilateral mOFC, rACC, PCC, OC as well as right-hemispheric VS and IFG/MFG, and left SFG, in accord with previous reports (Barberini et al., 2012). For instance, the mOFC plays specific roles in the consummatory phase of win but not loss processing (Chen et al., 2022a). The mOFC, ACC, and VS encode both the subjective value of drug-related cues and actions to procure the drug (Kable and Glimcher, 2007; McBride et al., 2006; Walton et al., 2004). These regions are engaged in responses to food and money (Delgado, 2007; Levy and Glimcher, 2012) as well as social interaction (Izuma et al., 2008), consistent with the findings of expectancy to increase social and physical pleasure in the approach biotype of drinkers. These brain regions are anatomically inter-connected to support appetitive cue responses (Makris et al., 2016; Ohtani et al., 2014a; Ohtani et al., 2014b). In support, patients with AUD relative to healthy individuals showed altered functional connectivity between the VS and medial prefrontal cortex (mPFC) in response to wins vs. losses, and stronger VS-mPFC connectivity was associated with higher drinking frequency and alcohol craving (Forbes et al., 2014; Park et al., 2010). Rodent studies also showed the dynamic activity of the VS circuit in encoding social rewards (Gunaydin et al., 2014).

The aversive circuit involved the AI, caudate and MCC, consistent with earlier findings of these regional roles in punishment-based avoidance learning (Knutson et al., 2014; Rolls, 2019). The involvement of AI in aversive responses is compatible with the insula in processing salient stimuli and interoceptive signals (Menon and Uddin, 2010; Naqvi and Bechara, 2010). The interoceptive circuit integrates multiple sources of information to support the experience of craving as well as behavioral control and emotion regulation (Craig, 2003). A human brain lesion study demonstrated that the AI was engaged in learning the negative values of loss cues and the dorsal striatum was involved in associative and motor aspects of decision-making to avoid the worst (Palminteri et al., 2012). In a recent study of Human Connectome Project, stronger neural responses in the caudate to punishment were associated with more severe alcohol use severity (Li et al., 2022). As a node in the cingulo-fronto-parietal network, the MCC is involved in avoidance, fear, and pain processing (Rolls, 2019). We recently reported that midcingulate cortical activations interrelated chronic craving and physiological responses to negative emotions in individuals with cocaine use disorder (Zhornitsky et al., 2021). These earlier findings are consistent with aversive cue reactivity of the AI, caudate, and MCC.

### 4.2 Clinical characteristics of the biotypes

As compared to the approach biotype, the avoidance biotype of drinkers showed markedly higher levels of punishment sensitivity, suggesting that these drinkers are more prone to respond with avoidance behaviors and likely engage in alcohol use to alleviate emotional distress. Earlier studies have implicated the insula, caudate, and MCC in negative emotions and avoidance behavior (Carretie et al., 2009; Rolls, 2019; Singer et al., 2009). For instance, sensitivity to punishment as assessed by Behavioral Inhibition System scale was associated with higher activation in the insula and dorsal striatum during avoidance anticipation (Kim et al., 2015). Thus, these trait differences are consistent with AI, caudate, and MCC cue reactivities as the neural markers of avoidance biotype. In contrast, drinkers of the approach vs. avoidance biotype showed higher levels of sensation seeking and expectancy for social and physical pleasure, suggesting that individuals of the approach biotype drink to enhance positive emotional state. The appetitive circuit that we identified involves regions implicated in sensation seeking (Chen et al., 2022b). For instance, the UPPS sensation seeking was positively associated with responses in the frontostriatal network to alcohol cues in heavy drinkers (Burnette et al., 2019). A previous study showed that craving ratings during cue exposure were positively correlated with AEQ social and physical pleasure (Carter, 2006) and stronger responses of the fusiform gyrus to positive vs. neutral words was correlated with higher level of expectancy for social and physical pleasure (Brislin et al., 2020).

Notably, the great majority of drinkers demonstrated a mixed psychological state during cue exposure, who showed intermediate levels of punishment sensitivity and sensation seeking relative to approach and avoidance biotypes. One important consideration is that although the MIDT involves win and loss trials, at stake is money that can be won or not lost. Thus, it is possible that participants were expecting to respond quickly so they will not lose, effectively winning, during these “loss trials;” the psychological processes involved may not be truly distinct from those of win trials. Thus, as encouraging as the results here, it is likely that to truly differentiate appetitive and aversive circuit activities, one would have to employ stimuli/outcomes of categorical differences, for instance, by replacing monetary loss with the delivery of an electric shock. A second consideration is that the meta-analyses identified ROIs across participants. It is likely that individuals vary in the appetitive and aversive processes and analyses of within-subject incentive and cue reactivity would reveal more robust findings. This would require experiments querying both cue and valenced motivational reactivity within the same participants.

### 4.3 Limitations of the study, other considerations, and conclusions

A few limitations should be considered. Firstly, only 20% of the drinkers in the current study reported an AUDIT score > 14, suggesting that the sample comprised largely non-dependent drinkers. Thus, the findings should be considered as specific to this population of social drinkers and drinkers with mild to severe alcohol use severity. Secondly, we did not examine sex differences because of the small sample size. Prior evidence showed that males drink more often in anticipation of positive emotions (Nolen-Hoeksema, 2004), whereas females are more likely to be motivated to drink and avoid aversive emotional states (Mooney et al., 1987). Indeed, we noticed that more drinkers in the approach biotype were male, although the sex composition was not significantly different from the avoidance biotype, likely due to the small sample size. It would be of tremendous interest to investigate how sex influences biotyping of drinkers.

To conclude, we characterized individual differences in the neural mechanisms of alcohol craving with respect to appetitive, aversive, or mixed nature of the psychological state. By biotyping alcohol drinkers according to these markers, we identified individual differences in the clinical characteristics and potentially the etiological processes of alcohol misuse.

## Acknowledgements

This study is supported by NIH grant DA051922. The funding agencies are otherwise not responsible for the design of the study, data collection or analysis, or in the decision to publish these results.

## Data/code availability statement

The data and codes will be shared on request to the corresponding author.

## References

Acar, F., Seurinck, R., Eickhoff, S.B., Moerkerke, B., 2018. Assessing robustness against potential publication bias in Activation Likelihood Estimation (ALE) meta-analyses for fMRI. PLoS One 13, e0208177.

American Psychiatric Association, A., 2013. Diagnostic and statistical manual of mental disorders (DSM-5®). American Psychiatric Pub.

Babor, T.F., Higgins-Biddle, J.C., Saunders, J.B., Monteiro, M.G., 2001. The alcohol use disorders identification test. World Health Organization Geneva.

Barberini, C.L., Morrison, S.E., Saez, A., Lau, B., Salzman, C.D., 2012. Complexity and competition in appetitive and aversive neural circuits. Front Neurosci 6, 170.

Bjork, J.M., Smith, A.R., Chen, G., Hommer, D.W., 2010. Adolescents, adults and rewards: comparing motivational neurocircuitry recruitment using fMRI. PLoS One 5, e11440.

Bonson, K.R., Grant, S.J., Contoreggi, C.S., Links, J.M., Metcalfe, J., Weyl, H.L., Kurian, V., Ernst, M., London, E.D., 2002. Neural systems and cue-induced cocaine craving. Neuropsychopharmacology 26, 376–386.

Brislin, S.J., Hardee, J.E., Martz, M.E., Cope, L.M., Weigard, A., Zucker, R.A., Heitzeg, M.M., 2020. Alcohol expectancies mediate the association between the neural response to emotional words and alcohol consumption. Drug Alcohol Depend 209, 107882.

Brody, A.L., Mandelkern, M.A., London, E.D., Childress, A.R., Lee, G.S., Bota, R.G., Ho, M.L., Saxena, S., Baxter, L.R., Madsen, D., 2002. Brain metabolic changes during cigarette craving. Archives of general psychiatry 59, 1162–1172.

Brown, S.A., Christiansen, B.A., Goldman, M.S., 1987. The Alcohol Expectancy Questionnaire: an instrument for the assessment of adolescent and adult alcohol expectancies. Journal of studies on alcohol 48, 483–491.

Burnette, E.M., Grodin, E.N., Lim, A.C., MacKillop, J., Karno, M.P., Ray, L.A., 2019. Association between impulsivity and neural activation to alcohol cues in heavy drinkers. Psychiatry Res Neuroimaging 293, 110986.

Carretie, L., Rios, M., de la Gandara, B.S., Tapia, M., Albert, J., Lopez-Martin, S., Alvarez-Linera, J., 2009. The striatum beyond reward: caudate responds intensely to unpleasant pictures. Neuroscience 164, 1615–1622.

Carter, A.C., 2006. Cue reactivity and the role of social alcohol expectancies in the college-aged drinking population.

Chen, Y., Chaudhary, S., Chiang-shan, R.L., 2022a. Shared and distinct neural activity during anticipation and outcome of win and loss: A meta-analysis of the monetary incentive delay task. bioRxiv.

Chen, Y., Ide, J.S., Li, C.S., Chaudhary, S., Le, T.M., Wang, W., Zhornitsky, S., Zhang, S., Li, C.S.R., 2022b. Gray matter volumetric correlates of dimensional impulsivity traits in children: Sex differences and heritability. Human brain mapping 43.

Clark, D.B., Thatcher, D.L., Tapert, S.F., 2008. Alcohol, psychological dysregulation, and adolescent brain development. Alcohol Clin Exp Res 32, 375–385.

Colledge, F., Ludyga, S., Mücke, M., Pühse, U., Gerber, M., 2018. The effects of an acute bout of exercise on neural activity in alcohol and cocaine craving: study protocol for a randomised controlled trial. Trials 19, 1–11.

Craig, A.D., 2003. Interoception: the sense of the physiological condition of the body. Current opinion in neurobiology 13, 500–505.

Delgado, M.R., 2007. Reward-related responses in the human striatum. Ann N Y Acad Sci 1104, 70–88.

Dhingra, I., Zhang, S., Zhornitsky, S., Le, T.M., Wang, W., Chao, H.H., Levy, I., Li, C.R., 2020. The effects of age on reward magnitude processing in the monetary incentive delay task. Neuroimage 207, 116368.

Dhingra, I., Zhang, S., Zhornitsky, S., Wang, W., Le, T.M., Li, C.R., 2021. Sex differences in neural responses to reward and the influences of individual reward and punishment sensitivity. BMC Neurosci 22, 12.

Eickhoff, S.B., Bzdok, D., Laird, A.R., Kurth, F., Fox, P.T., 2012. Activation likelihood estimation meta-analysis revisited. Neuroimage 59, 2349–2361.

Eickhoff, S.B., Bzdok, D., Laird, A.R., Roski, C., Caspers, S., Zilles, K., Fox, P.T., 2011. Co-activation patterns distinguish cortical modules, their connectivity and functional differentiation. Neuroimage 57, 938–949.

Eickhoff, S.B., Laird, A.R., Grefkes, C., Wang, L.E., Zilles, K., Fox, P.T., 2009. Coordinate-based activation likelihood estimation meta-analysis of neuroimaging data: a random-effects approach based on empirical estimates of spatial uncertainty. Hum Brain Mapp 30, 2907–2926.

Forbes, E.E., Rodriguez, E.E., Musselman, S., Narendran, R., 2014. Prefrontal response and frontostriatal functional connectivity to monetary reward in abstinent alcohol-dependent young adults. PLoS One 9, e94640.

Fox, H.C., Bergquist, K.L., Hong, K.I., Sinha, R., 2007. Stress-induced and alcohol cue-induced craving in recently abstinent alcohol-dependent individuals. Alcoholism: Clinical and Experimental Research 31, 395–403.

Frances, A., First, M.B., Pincus, H.A., 1995. DSM-IV guidebook. American Psychiatric Association.

Galandra, C., Basso, G., Cappa, S., Canessa, N., 2018. The alcoholic brain: neural bases of impaired reward-based decision-making in alcohol use disorders. Neurol Sci 39, 423–435.

George, O., Koob, G.F., 2022. Individual differences in the neuropsychopathology of addiction. Dialogues in clinical neuroscience.

Gunaydin, L.A., Grosenick, L., Finkelstein, J.C., Kauvar, I.V., Fenno, L.E., Adhikari, A., Lammel, S., Mirzabekov, J.J., Airan, R.D., Zalocusky, K.A., Tye, K.M., Anikeeva, P., Malenka, R.C., Deisseroth, K., 2014. Natural neural projection dynamics underlying social behavior. Cell 157, 1535–1551.

Heatherton, T.F., Kozlowski, L.T., Frecker, R.C., Fagerstrom, K.O., 1991. The Fagerström test for nicotine dependence: a revision of the Fagerstrom Tolerance Questionnaire. British journal of addiction 86, 1119–1127.

Hill-Bowen, L.D., Riedel, M.C., Poudel, R., Salo, T., Flannery, J.S., Camilleri, J.A., Eickhoff, S.B., Laird, A.R., Sutherland, M.T., 2021. The cue-reactivity paradigm: An ensemble of networks driving attention and cognition when viewing drug and natural reward-related stimuli. Neuroscience & Biobehavioral Reviews 130, 201–213.

Izuma, K., Saito, D.N., Sadato, N., 2008. Processing of social and monetary rewards in the human striatum. Neuron 58, 284–294.

Jasinska, A.J., Stein, E.A., Kaiser, J., Naumer, M.J., Yalachkov, Y., 2014. Factors modulating neural reactivity to drug cues in addiction: a survey of human neuroimaging studies. Neuroscience & Biobehavioral Reviews 38, 1–16.

Kable, J.W., Glimcher, P.W., 2007. The neural correlates of subjective value during intertemporal choice. Nat Neurosci 10, 1625–1633.

Kilts, C.D., Schweitzer, J.B., Quinn, C.K., Gross, R.E., Faber, T.L., Muhammad, F., Ely, T.D., Hoffman, J.M., Drexler, K.P., 2001. Neural activity related to drug craving in cocaine addiction. Archives of general psychiatry 58, 334–341.

Kim, S.H., Yoon, H., Kim, H., Hamann, S., 2015. Individual differences in sensitivity to reward and punishment and neural activity during reward and avoidance learning. Soc Cogn Affect Neurosci 10, 1219–1227.

King, A.C., McNamara, P.J., Hasin, D.S., Cao, D., 2014. Alcohol challenge responses predict future alcohol use disorder symptoms: a 6-year prospective study. Biological Psychiatry 75, 798–806.

Knutson, B., Katovich, K., Suri, G., 2014. Inferring affect from fMRI data. Trends Cogn Sci 18, 422–428.

Kuhn, S., Gallinat, J., 2011. Common biology of craving across legal and illegal drugs - a quantitative meta-analysis of cue-reactivity brain response. Eur J Neurosci 33, 1318–1326.

Lancaster, J.L., Tordesillas-Gutierrez, D., Martinez, M., Salinas, F., Evans, A., Zilles, K., Mazziotta, J.C., Fox, P.T., 2007. Bias between MNI and Talairach coordinates analyzed using the ICBM-152 brain template. Hum Brain Mapp 28, 1194–1205.

Le, T.M., Malone, T., Li, C.-S.R., 2022. Positive alcohol expectancy and resting-state functional connectivity of the insula in problem drinking. Drug and alcohol dependence 231, 109248.

Le, T.M., Zhornitsky, S., Zhang, S., Li, C.R., 2020. Pain and reward circuits antagonistically modulate alcohol expectancy to regulate drinking. Transl Psychiatry 10, 220.

Levy, D.J., Glimcher, P.W., 2012. The root of all value: a neural common currency for choice. Curr Opin Neurobiol 22, 1027–1038.

Li, G., Chen, Y., Chaudhary, S., Tang, X., Li, C.R., 2022. Loss and frontal striatal reactivities characterize alcohol use severity and rule-breaking behavior in young adult drinkers. Biol Psychiatry Cogn Neurosci Neuroimaging.

Limbrick-Oldfield, E.H., Mick, I., Cocks, R., McGonigle, J., Sharman, S., Goldstone, A.P., Stokes, P., Waldman, A., Erritzoe, D., Bowden-Jones, H., 2017. Neural substrates of cue reactivity and craving in gambling disorder. Translational psychiatry 7, e992–e992.

Makris, N., Rathi, Y., Mouradian, P., Bonmassar, G., Papadimitriou, G., Ing, W.I., Yeterian, E.H., Kubicki, M., Eskandar, E.N., Wald, L.L., Fan, Q., Nummenmaa, A., Widge, A.S., Dougherty, D.D., 2016. Variability and anatomical specificity of the orbitofrontothalamic fibers of passage in the ventral capsule/ventral striatum (VC/VS): precision care for patient-specific tractography-guided targeting of deep brain stimulation (DBS) in obsessive compulsive disorder (OCD). Brain Imaging Behav 10, 1054–1067.

McBride, D., Barrett, S.P., Kelly, J.T., Aw, A., Dagher, A., 2006. Effects of expectancy and abstinence on the neural response to smoking cues in cigarette smokers: an fMRI study. Neuropsychopharmacology 31, 2728–2738.

McClernon, F.J., Hiott, F.B., Huettel, S.A., Rose, J.E., 2005. Abstinence-induced changes in self-report craving correlate with event-related FMRI responses to smoking cues. Neuropsychopharmacology 30, 1940–1947.

Menon, V., Uddin, L.Q., 2010. Saliency, switching, attention and control: a network model of insula function. Brain Struct Funct 214, 655–667.

Mooney, D.K., Fromme, K., Kivlahan, D.R., Marlatt, G.A., 1987. Correlates of alcohol consumption: Sex, age, and expectancies relate differentially to quantity and frequency. Addictive behaviors 12, 235–240.

Muller, V.I., Cieslik, E.C., Laird, A.R., Fox, P.T., Radua, J., Mataix-Cols, D., Tench, C.R., Yarkoni, T., Nichols, T.E., Turkeltaub, P.E., Wager, T.D., Eickhoff, S.B., 2018. Ten simple rules for neuroimaging meta-analysis. Neurosci Biobehav Rev 84, 151–161.

Myrick, H., Anton, R.F., Li, X., Henderson, S., Drobes, D., Voronin, K., George, M.S., 2004. Differential brain activity in alcoholics and social drinkers to alcohol cues: relationship to craving. Neuropsychopharmacology 29, 393–402.

Naqvi, N.H., Bechara, A., 2010. The insula and drug addiction: an interoceptive view of pleasure, urges, and decision-making. Brain Struct Funct 214, 435–450.

Nolen-Hoeksema, S., 2004. Gender differences in risk factors and consequences for alcohol use and problems. Clin Psychol Rev 24, 981–1010.

Noori, H.R., Cosa Linan, A., Spanagel, R., 2016. Largely overlapping neuronal substrates of reactivity to drug, gambling, food and sexual cues: A comprehensive meta-analysis. Eur Neuropsychopharmacol 26, 1419–1430.

Ohtani, T., Bouix, S., Hosokawa, T., Saito, Y., Eckbo, R., Ballinger, T., Rausch, A., Melonakos, E., Kubicki, M., 2014a. Abnormalities in white matter connections between orbitofrontal cortex and anterior cingulate cortex and their associations with negative symptoms in schizophrenia: a DTI study. Schizophr Res 157, 190–197.

Ohtani, T., Nestor, P.G., Bouix, S., Saito, Y., Hosokawa, T., Kubicki, M., 2014b. Medial frontal white and gray matter contributions to general intelligence. PLoS One 9, e112691.

Oldham, S., Murawski, C., Fornito, A., Youssef, G., Yucel, M., Lorenzetti, V., 2018. The anticipation and outcome phases of reward and loss processing: A neuroimaging meta-analysis of the monetary incentive delay task. Hum Brain Mapp 39, 3398–3418.

Palminteri, S., Justo, D., Jauffret, C., Pavlicek, B., Dauta, A., Delmaire, C., Czernecki, V., Karachi, C., Capelle, L., Durr, A., Pessiglione, M., 2012. Critical roles for anterior insula and dorsal striatum in punishment-based avoidance learning. Neuron 76, 998–1009.

Park, S.Q., Kahnt, T., Beck, A., Cohen, M.X., Dolan, R.J., Wrase, J., Heinz, A., 2010. Prefrontal cortex fails to learn from reward prediction errors in alcohol dependence. J Neurosci 30, 7749–7753.

Pessiglione, M., Delgado, M.R., 2015. The good, the bad and the brain: Neural correlates of appetitive and aversive values underlying decision making. Curr Opin Behav Sci 5, 78–84.

Rolls, E.T., 2019. The cingulate cortex and limbic systems for emotion, action, and memory. Brain Struct Funct 224, 3001–3018.

Rosenthal, R., 1979. The file drawer problem and tolerance for null results. Psychological bulletin 86.

Schacht, J.P., Anton, R.F., Myrick, H., 2013. Functional neuroimaging studies of alcohol cue reactivity: a quantitative meta-analysis and systematic review. Addict Biol 18, 121–133.

Sell, L., Morris, J., Bearn, J., Frackowiak, R., Friston, K., Dolan, R.J., 1999. Activation of reward circuitry in human opiate addicts. European Journal of Neuroscience 11, 1042–1048.

Singer, T., Critchley, H.D., Preuschoff, K., 2009. A common role of insula in feelings, empathy and uncertainty. Trends Cogn Sci 13, 334–340.

Sinha, R., Fox, H.C., Hong, K.A., Bergquist, K., Bhagwagar, Z., Siedlarz, K.M., 2009. Enhanced negative emotion and alcohol craving, and altered physiological responses following stress and cue exposure in alcohol dependent individuals. Neuropsychopharmacology 34, 1198–1208.

Tiffany, S.T., 1990. A cognitive model of drug urges and drug-use behavior: role of automatic and nonautomatic processes. Psychological review 97, 147.

Torrubia, R., Avila, C., Moltó, J., Caseras, X., 2001. The Sensitivity to Punishment and Sensitivity to Reward Questionnaire (SPSRQ) as a measure of Gray’s anxiety and impulsivity dimensions. Personality and Individual Differences 31, 837–862.

Trotzke, P., Starcke, K., Pedersen, A., Brand, M., 2021. Dorsal and ventral striatum activity in individuals with buying-shopping disorder during cue-exposure: A functional magnetic resonance imaging study. Addiction Biology 26, e13073.

Turkeltaub, P.E., Eickhoff, S.B., Laird, A.R., Fox, M., Wiener, M., Fox, P., 2012. Minimizing within-experiment and within-group effects in activation likelihood estimation meta-analyses. Human brain mapping 33, 1–13.

Vafaie, N., Kober, H., 2022. Association of Drug Cues and Craving With Drug Use and Relapse: A Systematic Review and Meta-analysis. JAMA Psychiatry 79, 641–650.

Walton, M.E., Devlin, J.T., Rushworth, M.F., 2004. Interactions between decision making and performance monitoring within prefrontal cortex. Nat Neurosci 7, 1259–1265.

Wang, G.-J., Volkow, N.D., Fowler, J.S., Cervany, P., Hitzemann, R.J., Pappas, N.R., Wong, C.T., Felder, C., 1999. Regional brain metabolic activation during craving elicited by recall of previous drug experiences. Life sciences 64, 775–784.

Wang, W., Zhornitsky, S., Le, T.M., Zhang, S., Li, C.R., 2020. Heart Rate Variability, Cue-Evoked Ventromedial Prefrontal Cortical Response, and Problem Alcohol Use in Adult Drinkers. Biol Psychiatry Cogn Neurosci Neuroimaging 5, 619–628.

Whiteside, S.P., Lynam, D.R., Miller, J.D., Reynolds, S.K., 2005. Validation of the UPPS impulsive behaviour scale: a four-factor model of impulsivity. European Journal of Personality 19, 559–574.

Wilson, R.P., Colizzi, M., Bossong, M.G., Allen, P., Kempton, M., Mtac, Bhattacharyya, S., 2018. The Neural Substrate of Reward Anticipation in Health: A Meta-Analysis of fMRI Findings in the Monetary Incentive Delay Task. Neuropsychol Rev 28, 496–506.

Zeng, J., Yu, S., Cao, H., Su, Y., Dong, Z., Yang, X., 2021. Neurobiological correlates of cue-reactivity in alcohol-use disorders: A voxel-wise meta-analysis of fMRI studies. Neurosci Biobehav Rev 128, 294–310.

Zhang, S., Zhornitsky, S., Wang, W., Le, T.M., Dhingra, I., Chen, Y., Li, C.s.R., 2021. Resting state hypothalamic and dorsomedial prefrontal cortical connectivity of the periaqueductal gray in cocaine addiction. Addiction Biology 26, e12989.

Zhornitsky, S., Le, T.M., Wang, W., Dhingra, I., Chen, Y., Chiang-shan, R.L., Zhang, S., 2021. Midcingulate Cortical Activations Interrelate Chronic Craving and Physiological Responses to Negative Emotions in Cocaine Addiction. Biological Psychiatry Global Open Science 1, 37–47.

Zhornitsky, S., Zhang, S., Ide, J.S., Chao, H.H., Wang, W., Le, T.M., Leeman, R.F., Bi, J., Krystal, J.H., Chiang-shan, R.L., 2019. Alcohol expectancy and cerebral responses to cue-elicited craving in adult nondependent drinkers. Biological psychiatry: cognitive neuroscience and neuroimaging 4, 493–504.

