## Supplement for "Appetitive and aversive cue reactivities differentiate biotypes of alcohol drinkers"

**Supplementary Table S1.** A list of 53 studies included in the meta-analysis of cue-elicited reactivity to drug/substance

| *No.* | *Study* | *n* | *Substance user group* | *Contrast* |
|---|---|---|---|---|
| 1 | Al-Khalil et al. (2021) | 149 | alcohol | alcohol vs. neutral |
| 2 | Allenby et al. (2020) | 75 | nicotine | smoking vs. neutral |
| 3 | Bach et al. (2015) | 81 | alcohol | alcohol vs. neutral |
| 4 | Bi et al. (2017) | 35 | nicotine | smoking vs. neutral |
| 5 | Bourque et al. (2013) | 31 | nicotine | smoking vs. neutral |
| 6 | Bradstreet et al. (2014) | 30 | nicotine | smoking vs. neutral |
| 7 | Chen et al. (2018) | 41 | nicotine | e-cigarette vs. neutral |
| 8 | Courtney et al. (2018) | 53 | alcohol | alcohol vs. neutral |
| 9 | Courtney et al. (2016) | 23 | methamphetamine | methamphetamine vs. neutral |
| 10 | Cousijn et al. (2013) | 31 | cannabis | cannabis vs. neutral |
| 11 | David et al. (2007) | 12 | nicotine | smoking vs. neutral |
| 12 | David et al. (2005) | 14 | nicotine | smoking vs. neutral |
| 13 | Falcone et al. (2016) | 69 | nicotine | smoking vs. neutral |
| 14 | Filbey et al. (2016) | 53 | cannabis | cannabis vs. neutral |
| 15 | George et al. (2001) | 10 | alcohol | alcohol vs. neutral |
| 16 | Grodin et al. (2021) | 25 | alcohol | alcohol vs. neutral |
| 17 | Hanlon et al. (2012) | 26 | nicotine | smoking vs. neutral |
| 18 | Hanlon et al. (2018) | 156 | alcohol/cocaine/nicotine | [a]drug vs. neutral |
| 19 | Hassani-Abharian et al. (2015) | 25 | heroin | heroin vs. neutral |
| 20 | Haugg et al. (2022) | 32 | nicotine | smoking vs. neutral |
| 21 | Janes et al. (2009) | 13 | nicotine | smoking vs. neutral |
| 22 | Janes et al. (2012) | 24 | nicotine | smoking vs. neutral |
| 23 | Janes et al. (2015) | 17 | nicotine | smoking vs. neutral |
| 24 | Kaag et al. (2018) | 51 | cocaine | cocaine vs. neutral |
| 25 | Karl et al. (2021) | 15 | alcohol | alcohol vs. neutral |
| 26 | Karoly et al. (2019) | 41 | cannabis | cannabis vs. neutral |
| 27 | Kim et al. (2020) | 30 | nicotine | smoking vs. neutral |
| 28 | Kleinhans et al. (2020) | 25 | cannabis | cannabis vs. neutral |
| 29 | Kushnir et al. (2013) | 18 | nicotine | smoking vs. neutral |
| 30 | Le et al. (2020) | 71 | alcohol | alcohol vs. neutral |
| 31 | Li et al. (2020) | 24 | nicotine | smoking vs. neutral |
| 32 | Malcolm et al. (2016) | 9 | methamphetamine | methamphetamine vs. neutral |
| 33 | McClernon et al. (2007) | 13 | nicotine | smoking vs. neutral |
| 34 | McClernon et al. (2008) | 30 | nicotine | smoking vs. neutral |
| 35 | Mondino et al. (2018) | 29 | nicotine | smoking vs. neutral |
| 36 | (Moran-Santa Maria et al., 2015) | 58 | nicotine | smoking vs. neutral |
| 37 | Myrick et al. (2008) | 24 | alcohol | alcohol vs. neutral |
| 38 | Myrick et al. (2010) | 16 | alcohol | alcohol vs. neutral |
| 39 | Park et al. (2007) | 9 | alcohol | alcohol vs. neutral |
| 40 | Potvin et al. (2016) | 24 | nicotine | smoking vs. neutral |
| 41 | Ray et al. (2010) | 10 | alcohol/marijuana | [b]drug vs. neutral |
| 42 | Regier et al. (2017) | 40 | cocaine | cocaine vs. neutral |

| | | | | |
|---|---|---|---|---|
| 43 | Rubinstein et al. (2011) | 12 | nicotine | smoking vs. neutral |
| 44 | Schulte et al. (2019) | 15 | cocaine | cocaine vs. neutral |
| 45 | Shi et al. (2021) | 29 | opioid | opioid vs. neutral |
| 46 | Sjoerds et al. (2014) | 30 | alcohol | alcohol vs. neutral |
| 47 | Versace et al. (2011) | 35 | nicotine | smoking vs. neutral |
| 48 | Vollstadt-Klein et al. (2011) | 22 | nicotine | smoking vs. neutral |
| 49 | Vollstadt-Klein et al. (2010) | 31 | alcohol | alcohol vs. neutral |
| 50 | Walter et al. (2015) | 27 | heroin | heroin vs. neutral |
| 51 | Wang et al. (2021) | 44 | cocaine | cocaine vs. neutral |
| 52 | Zhang et al. (2020) | 23 | cocaine | cocaine vs. neutral |
| 53 | Zhornitsky et al. (2019) | 61 | alcohol | alcohol vs. neutral |

*Note:* [a]Stimuli include alcohol, cocaine, and smoking pictures; [b]Stimuli include cocaine and marijuana pictures.

**Supplementary Table S2.** ALE results for activations for drug > neutral cue in the cue craving task

| Cluster volumes ($mm^3$) | ALE | MNI Coordinates (mm) | | | Identified Region |
|---|---|---|---|---|---|
| | | x | y | z | |
| 12,896 | 0.05 | 6 | 52 | -4 | L dorsal anterior cingulate cortex |
| | 0.04 | -10 | 44 | -12 | L dorsal anterior cingulate cortex |
| | 0.03 | -2 | 38 | 4 | L ventral anterior cingulate cortex |
| | 0.03 | 0 | 36 | 10 | L ventral anterior cingulate cortex |
| | 0.02 | -2 | 54 | 16 | L medial frontal gyrus |
| 8,096 | 0.04 | -6 | -48 | 26 | L posterior cingulate cortex |
| | 0.04 | 6 | -46 | 24 | R posterior cingulate cortex |
| | 0.03 | -4 | -34 | 32 | L cingulate cortex |
| | 0.03 | -4 | -58 | 14 | L posterior cingulate cortex |
| | 0.02 | 2 | -16 | 36 | L ventral anterior cingulate cortex |
| | 0.02 | 0 | -20 | 32 | L cingulate cortex |
| | 0.02 | 0 | 2 | 30 | L ventral anterior cingulate cortex |
| 2,808 | 0.04 | -18 | 44 | 42 | L superior frontal gyrus |
| | 0.02 | -12 | 48 | 32 | L medial frontal gyrus |
| *Evaluation of publication bias with additional 275 null studies* | | | | | |
| 2,368 | 0.06 | -4 | 50 | -6 | L dorsal anterior cingulate cortex |

*Note:* The results were evaluated at voxel-level $p < 0.001$ in combination with cluster-level $p < 0.05$ family-wise error (FWE) corrected; L: Left; R: right.

**Supplementary Table 3** ALE results for activations for win and loss processing in the MIDT

| Cluster volumes ($mm^3$) | ALE | MNI Coordinates (mm) | | | Identified Region |
|---|---|---|---|---|---|
| | | x | y | z | |
| ***winAO*** | | | | | |
| 43,592 | 0.23 | 12 | 10 | -4 | R caudate head |
| | 0.18 | -10 | 8 | -4 | L caudate head |
| | 0.10 | 34 | 24 | -2 | R insula |
| | 0.07 | -30 | 26 | 4 | L insula |

| | | | | | |
|---|---|---|---|---|---|
| | 0.07 | 4 | -12 | 6 | R thalamus, medial dorsal nucleus |
| | 0.06 | -4 | -10 | 8 | L thalamus, medial dorsal nucleus |
| | 0.06 | -6 | -26 | -4 | L thalamus |
| | 0.06 | -8 | -16 | 8 | L thalamus, medial dorsal nucleus |
| | 0.05 | 8 | -24 | -4 | R thalamus |
| | 0.04 | 0 | -20 | -14 | L midbrain, red nucleus |
| | 0.03 | -14 | -2 | 14 | L caudate body |
| | 0.03 | 20 | -22 | -14 | R parahippocampal gyrus |
| | 0.03 | -18 | 0 | 18 | L caudate body |
| | 0.03 | 18 | 4 | 16 | R caudate body |
| 6,472 | 0.10 | 4 | 6 | 52 | R medial frontal gyrus |
| 4,864 | 0.07 | 24 | -90 | -4 | R middle occipital gyrus |
| | 0.06 | 16 | -88 | -4 | R lingual gyrus |
| | 0.04 | 34 | -90 | 6 | R middle occipital gyrus |
| | 0.03 | 6 | -80 | -2 | R lingual gyrus |
| 3,976 | 0.05 | -16 | -88 | -10 | L lingual gyrus |
| | 0.04 | -12 | -94 | 0 | L lingual gyrus |
| | 0.04 | -24 | -92 | 2 | L lingual gyrus |
| | 0.04 | -28 | -88 | -8 | L middle occipital gyrus |
| 3,208 | 0.08 | -38 | -16 | 52 | L precentral gyrus |
| 1,392 | 0.06 | 44 | -2 | 50 | R precentral gyrus |
| 880 | 0.04 | 2 | -38 | 36 | L cingulate gyrus |
| | 0.03 | 0 | -30 | 26 | L posterior cingulate gyrus |
| ***lossAO*** | | | | | |
| 30,248 | 0.09 | 12 | 10 | -2 | R caudate head |
| | 0.09 | -10 | 8 | -2 | L caudate head |
| | 0.04 | 34 | 26 | -2 | R insula |
| | 0.04 | -8 | -16 | 4 | L thalamus |
| | 0.04 | 12 | 0 | 14 | R caudate body |
| | 0.03 | 42 | 20 | -2 | R insula |
| | 0.03 | 0 | -28 | -4 | L midbrain, red nucleus |
| | 0.03 | 6 | -12 | 6 | R thalamus, medial dorsal nucleus |
| | 0.02 | 8 | -24 | -10 | R midbrain, red nucleus |
| 5,136 | 0.06 | 4 | 2 | 54 | R medial frontal gyrus |
| | 0.03 | 6 | 16 | 40 | R cingulate gyrus |
| 2,456 | 0.04 | -30 | 24 | -4 | L claustrum |
| | 0.03 | -40 | 12 | -2 | L insula |
| 1,912 | 0.04 | -16 | -90 | -8 | L lingual gyrus |
| | 0.03 | -12 | -94 | 0 | L lingual gyrus |
| 1,544 | 0.04 | 42 | -2 | 48 | R middle frontal gyrus |
| 1,480 | 0.03 | -40 | -14 | 54 | L precentral gyrus |
| 840 | 0.03 | -28 | -60 | 44 | L precuneus |

*Note:* The results were evaluated at voxel-level $p < 0.001$ in combination with cluster-level $p < 0.05$ family-wise error (FWE) corrected. Win AO: win anticipation and outcome; lossAO: loss anticipation and outcome; L: Left; R: right.

**Supplementary Table S4.** ALE results for subtraction analyses (winAO > lossAO and lossAO > win AO), evaluated at $p < 0.001$ or $p < 0.05$ with a minimal cluster size of 100 $mm^3$

| Cluster volumes ($mm^3$) | *Z* | MNI Coordinates (mm) | | | Identified Region |
|---|---|---|---|---|---|
| | | x | y | z | |
| ***p < 0.001*** | | | | | |
| *winAO > lossAO* | | | | | |
| 520 | 3.16 | 2.9 | -36 | 31.4 | R cingulate gyrus |
| | 3.24 | -2.3 | -40 | 32.6 | L cingulate gyrus |
| | 3.09 | -2 | -36 | 32 | L cingulate gyrus |
| *lossAO > winAO* | | | | | |
| Nil | | | | | |
| ***p < 0.05*** | | | | | |
| *winAO > lossAO* | | | | | |
| 4,024 | 2.77 | 12 | 10 | -10 | R caudate head |
| | 2.56 | 0 | 14 | -10 | L anterior cingulate |
| | 2.47 | -2 | 14 | -14 | L anterior cingulate |
| | 2.22 | 2 | 12 | -4 | L caudate head |
| 2,800 | 3.72 | -1.8 | 46.5 | -9.2 | L anterior cingulate |
| 2,664 | 3.72 | 0.6 | -40.4 | 32.9 | L cingulate gyrus |
| | 3.54 | 2.6 | -35.3 | 32 | L cingulate gyrus |
| | 3.24 | 0 | -32 | 36 | L cingulate gyrus |
| 2,008 | 2.55 | 22 | -84 | 0 | R lingual gyrus |
| | 2.39 | 37.3 | -88.7 | -6.7 | R inferior occipital gyrus |
| | 2.11 | 36 | -84 | -12 | R fusiform gyrus |
| | 1.81 | 28 | -78 | -10 | R lingual gyrus |
| 1,848 | 3.09 | -30 | 40 | 32 | L middle frontal gyrus |
| | 3.04 | -32 | 40 | 30 | L superior frontal gyrus |
| | 2.79 | -38.7 | 39 | 25 | L middle frontal gyrus |
| | 2.77 | -42 | 44 | 26 | L superior frontal gyrus |
| | 2.27 | -42 | 48 | 20 | L middle frontal gyrus |
| 1,776 | 3.35 | -36 | -84 | 4 | L inferior occipital gyrus |
| | 3.24 | -32 | -84 | 4 | L middle occipital gyrus |
| 1,592 | 3.35 | 34 | 42 | 32 | R middle frontal gyrus |
| | 2.55 | 38 | 52 | 28 | R middle frontal gyrus |
| 1,576 | 3.35 | 10 | -68 | 4 | R lingual gyrus |
| 1,256 | 2.85 | 12 | -24 | 6 | R thalamus, pulvinar |
| | 2.82 | 12 | -22 | 10 | R thalamus, medial dorsal nucleus |
| | 2.73 | 20 | -14 | 8 | R thalamus, ventral lateral nucleus |
| | 2.46 | 14 | -16 | 14 | R thalamus, lateral dorsal nucleus |
| 1,064 | 2.85 | -40 | -49 | -30 | L cerebellum |
| | 2.55 | -40 | -56 | -30 | L cerebellum |
| | 2.42 | -38 | -50 | -36 | L cerebellum |
| 936 | 2.69 | 33 | -68 | 36 | R precuneus |

| | | | | | |
|---|---|---|---|---|---|
| 600 | 2.44 | -8 | -14 | -14 | L midbrain, substania nigra |
| | 2.42 | -2 | -18 | -12 | L midbrain, red nucleus |
| | 1.85 | -12 | -12 | -12 | L midbrain, subthalamic nucleus |
| 504 | 2.24 | -26 | 10 | 0 | L putamen |
| | 2.11 | -20 | 16 | 4 | L putamen |
| | 2.67 | 0 | 8 | 32 | L cingulate gyrus |
| 504 | 2.65 | 28 | -22 | -12 | R parahippocampal gyrus |
| 496 | 2.56 | 28 | -26 | -20 | R parahippocampal gyrus |
| | 2.51 | 28 | -26 | -14 | R parahippocampal gyrus |
| | 2.41 | -22 | 0 | 16 | L putamen |
| 440 | 2.32 | 22 | -42 | -22 | R cerebellum |
| 376 | 1.93 | 48.7 | 33.3 | 16 | R middle frontal gyrus |
| 376 | 2.00 | -20 | 34 | 46 | L superior frontal gyrus |
| 344 | 1.98 | -16 | 30 | 48 | L superior frontal gyrus |
| | 2.18 | -50 | 18 | 26 | L middle frontal gyrus |
| 296 | 2.33 | 36 | -36 | -14 | R parahippocampal gyrus |
| 272 | 2.32 | 36 | -31 | -14 | R hippocampus |
| | 2.13 | 40 | -32 | -12 | R hippocampus |
| | 2.05 | 14 | -66 | 40 | R precuneus |
| 264 | 1.77 | 22 | -72 | 40 | R cuneus |
| | 2.69 | 26 | -16 | 0 | R lateral globus pallidus |
| 208 | 2.65 | 24 | -12 | 0 | R lateral globus pallidus |
| | 2.00 | -28 | -12 | 4 | L putamen |
| 200 | 1.74 | -30 | -10 | -2 | L putamen |
| | 2.10 | -32 | 34 | 4 | L inferior frontal gyrus |
| 200 | 2.26 | -40 | -26 | 62 | L postcentral gyrus |
| 200 | 2.49 | -4 | 38 | 8 | L anterior cingulate |
| 184 | 2.04 | -6 | 8 | 54 | L medial frontal gyrus |
| 184 | 2.25 | 44 | -54 | -14 | R fusiform gyrus |
| 112 | 1.87 | -2 | 24 | 32 | L cingulate gyrus |
| 104 | 1.81 | -6 | 26 | 34 | L cingulate gyrus |
| *lossAO > winAO* | | | | | |
| 3,200 | 3.54 | -7 | 8 | 12 | L caudate body |
| | 2.59 | -8 | -6 | 16 | L thalamus |
| | 2.48 | -6 | -6 | -4 | L thalamus |
| 1,672 | 0.00 | 47 | 23 | 4 | R inferior frontal gyrus |
| | 3.35 | 46 | 16 | 0 | R insula |
| | 2.10 | 42 | 30 | 2 | R inferior frontal gyrus |
| 1,208 | 2.26 | 2 | 26 | 48 | L superior frontal gyrus |
| | 2.20 | 2 | 32 | 50 | L superior frontal gyrus |
| | 1.99 | 4 | 40 | 42 | R superior frontal gyrus |
| | 1.89 | 4 | 40 | 34 | R medial frontal gyrus |
| 992 | 2.47 | 12 | -2 | 14 | R caudate body |
| 904 | 2.47 | 17.3 | -4 | 70 | R superior frontal gyrus |

| | | | | | |
|---|---|---|---|---|---|
| | 2.41 | 17 | -5 | 64 | R medial frontal gyrus |
| | 2.34 | 20.7 | -10 | 63.3 | R medial frontal gyrus |
| | 2.32 | 20 | -14 | 70 | R precentral gyrus |
| 872 | 2.67 | -42 | 14 | 0 | L insula |
| 744 | 2.40 | -8 | -76 | 10 | L lingual gyrus |
| | 2.20 | -14 | -86 | 10 | L cuneus |
| | 2.16 | -14 | -88 | 14 | L cuneus |
| 584 | 2.60 | 10 | -4 | 48 | R cingulate gyrus |
| | 2.39 | 6 | -10 | 44 | R cingulate gyrus |
| 560 | 2.17 | 50 | -10 | 42 | R precentral gyrus |
| | 2.15 | 58 | -4 | 40 | R precentral gyrus |
| 536 | 2.35 | -12 | -76 | -25 | L cerebellum |
| | 2.33 | -10 | -72 | -24 | L cerebellum |
| 456 | 2.07 | -51 | -10 | 39 | L precentral gyrus |
| 384 | 2.30 | -8 | -8 | 64 | L medial frontal gyrus |
| 360 | 1.97 | -26 | -62 | 48 | L superior parietal lobule |
| | 1.97 | -30 | -60 | 50 | L superior parietal lobule |
| 320 | 2.32 | 60 | 18 | 12 | R inferior frontal gyrus |
| 296 | 1.98 | 30 | -66 | -2 | R lingual gyrus |
| | 1.74 | 34 | -66 | -8 | R cerebellum |
| 216 | 2.12 | 4 | -12 | 42 | R cingulate gyrus |
| 216 | 1.84 | 6 | 8 | 68 | R superior frontal gyrus |
| | 1.72 | 10 | 8 | 68 | R superior frontal gyrus |
| 192 | 2.18 | -10 | -18 | -2 | L thalamus, mammillary body |
| 112 | 1.80 | 30 | -60 | 12 | R posterior cingulate |
| 112 | 1.98 | 12 | 20 | 40 | R medial frontal gyrus |
| 104 | 1.98 | 22 | 2 | 4 | R putamen |

*Note:* Win AO: win anticipation and outcome; lossAO: loss anticipation and outcome; Nil: no significant findings; L: Left; R: right.

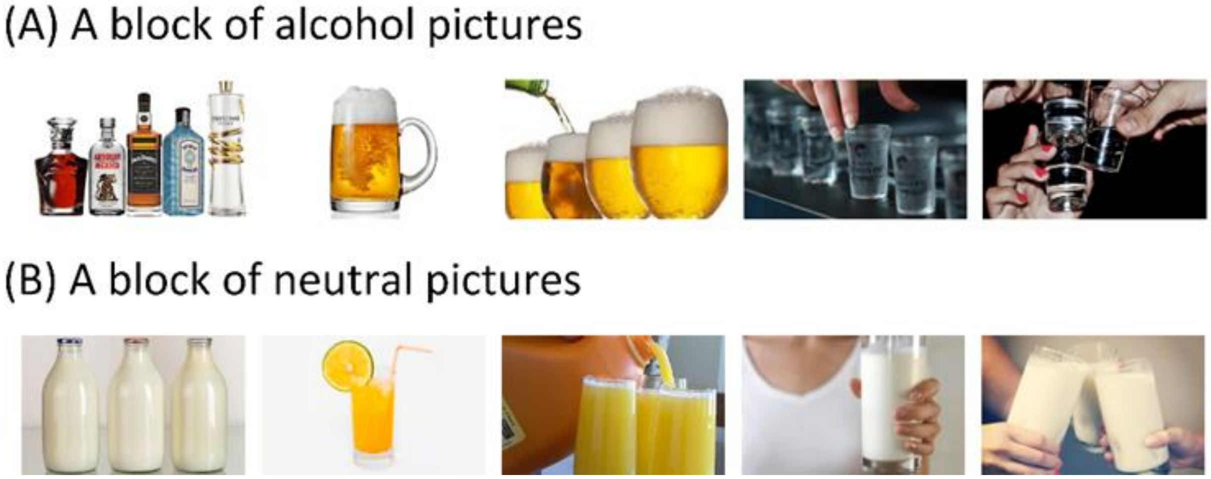

**Supplementary Figure S1.** Cue-induced alcohol craving task. Participants performed 2 runs, each containing 6 blocks of (**A**) alcohol cues and (**B**) neutral cues. Each picture was presented for 6 seconds. Blocks began with a 2-second fixation. Participants were instructed to report their alcohol craving at the end of every block. Each block lasted approximately 45 seconds, including the time for craving rating.

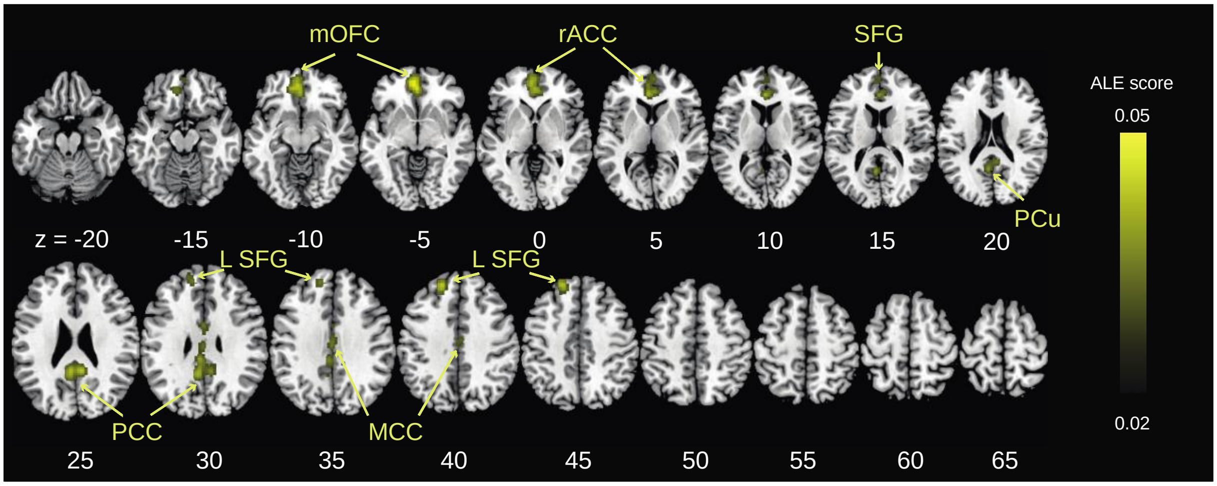


**Supplementary Figure S2.** Cue-elicited reactivity to drug. *Note:* The results were evaluated with a cluster-forming threshold of $p < 0.001$ uncorrected and a cluster-level threshold of $p < 0.05$ FWE corrected. Color bars represent ALE scores. L: left; MCC: mid-cingulate cortex; mOFC: medial orbitofrontal cortex; PCC: posterior cingulate cortex; PCu: precuneus; rACC: rostral anterior cingulate cortex; SFG: superior frontal cortex.

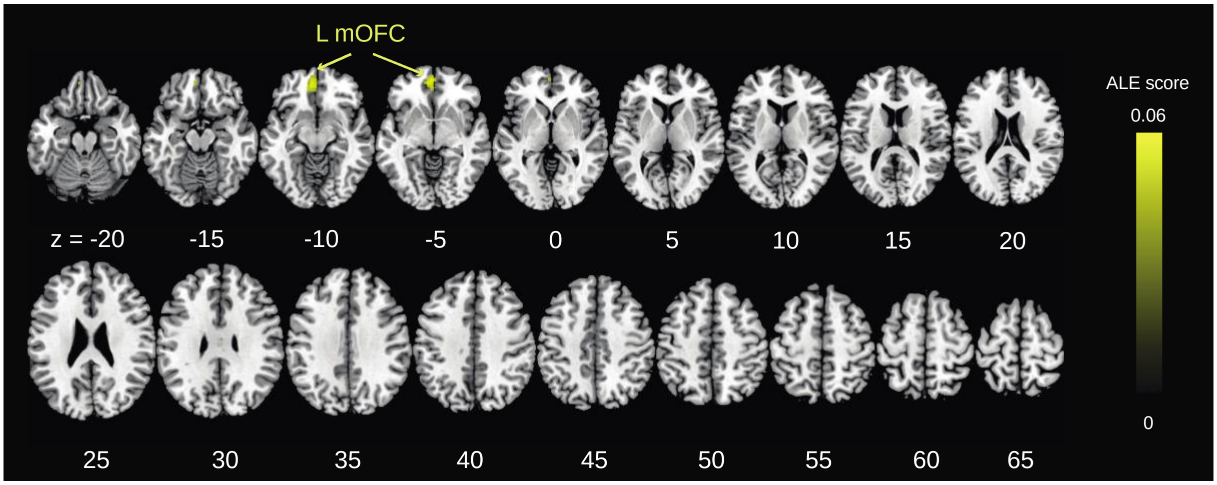


**Supplementary Figure S3.** Cue-elicited reactivity to drug, for the evaluation of publication bias with additional 275 null studies. *Note:* The results were evaluated with a cluster-forming threshold of $p < 0.001$ uncorrected and a cluster-level threshold of $p < 0.05$ FWE corrected. Color bars represent ALE scores.

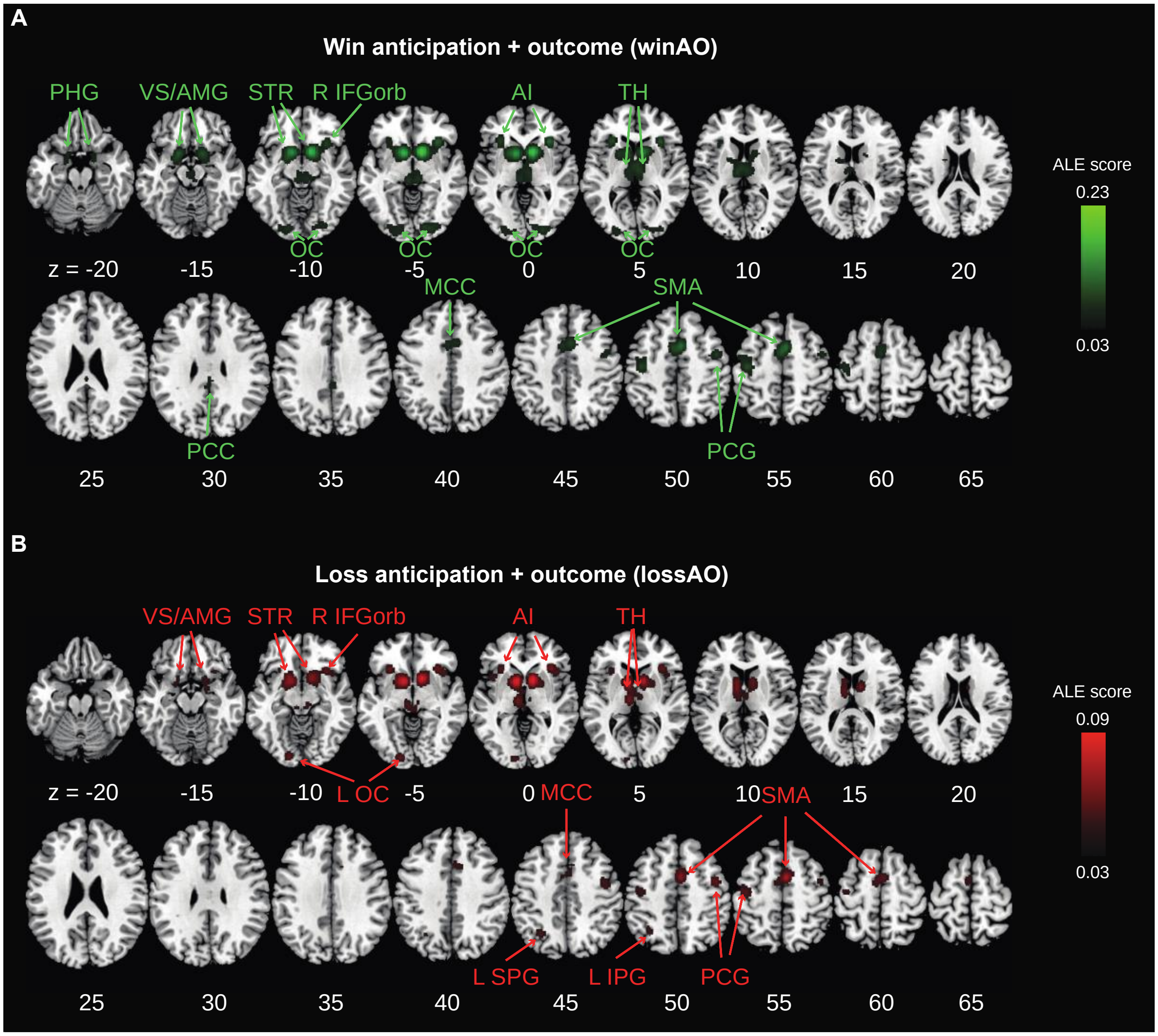


**Supplementary Figure S4.** ALE results for **(A)** Win anticipation and outcome (winAO) and **(B)** Loss anticipation and outcome (lossAO). *Note:* The results were evaluated with a cluster-forming threshold of $p < 0.001$ uncorrected and a cluster-level threshold of $p < 0.05$ FWE corrected. Color bars represent ALE scores. L: left; R: right; AI: anterior cingulate cortex; AMG: amygdala; IFGorb: inferior frontal gyrus pars orbitalis; IPG: inferior parietal gyrus; MCC: mid-cingulate cortex; OC: occipital cortex; PCC: posterior cingulate cortex; PCG: precentral gyrus; PHG: parahippocampal gyrus; SFG: superior frontal cortex; SMA: supplementary motor area; SPG: superior parietal gyrus; STR: striatum; TH: thalamus; VS: ventral striatum.

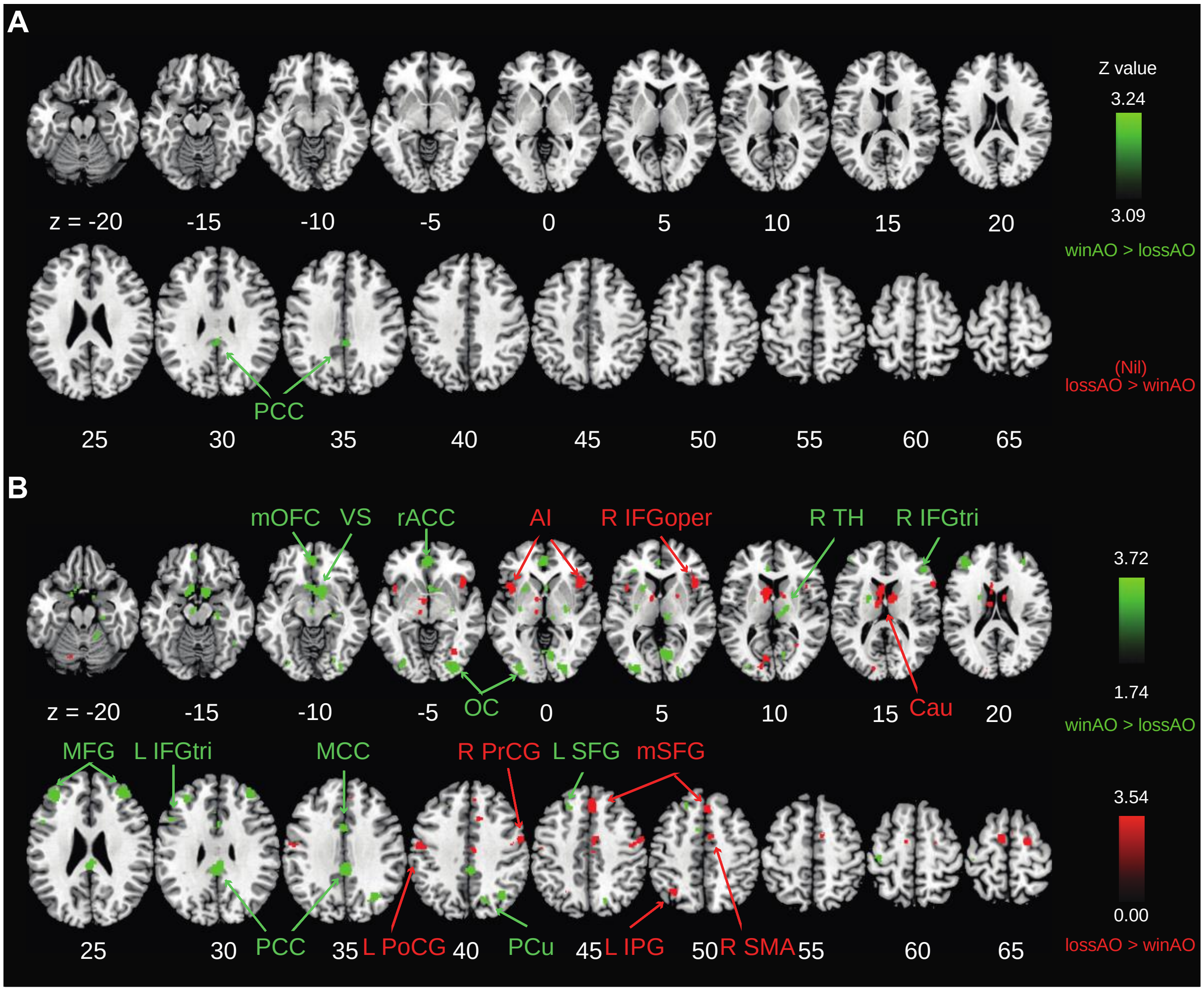


**Supplementary Figure S5.** ALE subtraction analyses of winAO > lossAO and lossAO > winAO with results evaluated at a significance level of (**A**) $p < 0.001$ and (**B**) $p < 0.05$ with a minimal cluster size of 100 mm$^3$. Color bars represent $Z$ values. winAO: win anticipation and outcome; lossAO: loss anticipation and outcome; Nil: no significant findings; PCC: posterior cingulate cortex.

### References

Al-Khalil, K., Vakamudi, K., Witkiewitz, K., Claus, E.D., 2021. Neural correlates of alcohol use disorder severity among nontreatment-seeking heavy drinkers: An examination of the incentive salience and negative emotionality domains of the alcohol and addiction research domain criteria. Alcohol Clin Exp Res 45, 1200-1214.

Allenby, C., Falcone, M., Wileyto, E.P., Cao, W., Bernardo, L., Ashare, R.L., Janes, A., Loughead, J., Lerman, C., 2020. Neural cue reactivity during acute abstinence predicts short-term smoking relapse. Addict Biol 25, e12733.

Bach, P., Vollsta Dt-Klein, S., Kirsch, M., Hoffmann, S., Jorde, A., Frank, J., Charlet, K., Beck, A., Heinz, A., Walter, H., Sommer, W.H., Spanagel, R., Rietschel, M., Kiefer, F., 2015. Increased mesolimbic cue-reactivity in carriers of the mu-opioid-receptor gene OPRM1 A118G polymorphism predicts drinking outcome: a functional imaging study in alcohol dependent subjects. Eur Neuropsychopharmacol 25, 1128-1135.

Bi, Y., Yuan, K., Yu, D., Wang, R., Li, M., Li, Y., Zhai, J., Lin, W., Tian, J., 2017. White matter integrity of central executive network correlates with enhanced brain reactivity to smoking cues. Hum Brain Mapp 38, 6239-6249.

Bourque, J., Mendrek, A., Dinh-Williams, L., Potvin, S., 2013. Neural circuitry of impulsivity in a cigarette craving paradigm. Front Psychiatry 4, 67.

Bradstreet, M.P., Higgins, S.T., McClernon, F.J., Kozink, R.V., Skelly, J.M., Washio, Y., Lopez, A.A., Parry, M.A., 2014. Examining the effects of initial smoking abstinence on response to smoking-related stimuli and response inhibition in a human laboratory model. Psychopharmacology (Berl) 231, 2145-2158.

Chen, Y., Fowler, C.H., Papa, V.B., Lepping, R.J., Brucks, M.G., Fox, A.T., Martin, L.E., 2018. Adolescents' behavioral and neural responses to e-cigarette advertising. Addict Biol 23, 761-771.

Courtney, A.L., Rapuano, K.M., Sargent, J.D., Heatherton, T.F., Kelley, W.M., 2018. Reward system activation in response to alcohol advertisements predicts college drinking. Journal of studies on alcohol and drugs 79, 29-38.

Courtney, K.E., Ghahremani, D.G., Ray, L.A., 2016. The Effects of Pharmacological Opioid Blockade on Neural Measures of Drug Cue-Reactivity in Humans. Neuropsychopharmacology 41, 2872-2881.

Cousijn, J., Goudriaan, A.E., Ridderinkhof, K.R., van den Brink, W., Veltman, D.J., Wiers, R.W., 2013. Neural responses associated with cue-reactivity in frequent cannabis users. Addict Biol 18, 570-580.

David, S.P., Munafo, M.R., Johansen-Berg, H., Mackillop, J., Sweet, L.H., Cohen, R.A., Niaura, R., Rogers, R.D., Matthews, P.M., Walton, R.T., 2007. Effects of Acute Nicotine Abstinence on Cue-elicited Ventral Striatum/Nucleus Accumbens Activation in Female Cigarette Smokers: A Functional Magnetic Resonance Imaging Study. Brain Imaging Behav 1, 43-57.

David, S.P., Munafo, M.R., Johansen-Berg, H., Smith, S.M., Rogers, R.D., Matthews, P.M., Walton, R.T., 2005. Ventral striatum/nucleus accumbens activation to smoking-related pictorial cues in smokers and nonsmokers: a functional magnetic resonance imaging study. Biol Psychiatry 58, 488-494.

Falcone, M., Cao, W., Bernardo, L., Tyndale, R.F., Loughead, J., Lerman, C., 2016. Brain Responses to Smoking Cues Differ Based on Nicotine Metabolism Rate. Biol Psychiatry 80, 190-197.

Filbey, F.M., Dunlop, J., Ketcherside, A., Baine, J., Rhinehardt, T., Kuhn, B., DeWitt, S., Alvi, T., 2016. fMRI study of neural sensitization to hedonic stimuli in long-term, daily cannabis users. Hum Brain Mapp 37, 3431-3443.

George, M.S., Anton, R.F., Bloomer, C., Teneback, C., Drobes, D.J., Lorberbaum, J.P., Nahas, Z., Vincent, D.J., 2001. Activation of prefrontal cortex and anterior thalamus in alcoholic subjects on exposure to alcohol-specific cues. Archives of general psychiatry 58, 345-352.

Grodin, E.N., Burnette, E.M., Green, R., Lim, A.C., Miotto, K., Ray, L.A., 2021. Combined varenicline and naltrexone attenuates alcohol cue-elicited activation in heavy drinking smokers. Drug Alcohol Depend 225, 108825.

Hanlon, C.A., Dowdle, L.T., Gibson, N.B., Li, X., Hamilton, S., Canterberry, M., Hoffman, M., 2018. Cortical substrates of cue-reactivity in multiple substance dependent populations: transdiagnostic relevance of the medial prefrontal cortex. Transl Psychiatry 8, 186.

Hanlon, C.A., Jones, E.M., Li, X., Hartwell, K.J., Brady, K.T., George, M.S., 2012. Individual variability in the locus of prefrontal craving for nicotine: implications for brain stimulation studies and treatments. Drug Alcohol Depend 125, 239-243.
Hassani-Abharian, P., Ganjgahi, H., Tabatabaei-Jafari, H., Oghabian, M.A., Mokri, A., Ekhtiari, H., 2015. Exploring neural correlates of different dimensions in drug craving self-reports among heroin dependents. Basic and clinical neuroscience.
Haugg, A., Manoliu, A., Sladky, R., Hulka, L.M., Kirschner, M., Bruhl, A.B., Seifritz, E., Quednow, B.B., Herdener, M., Scharnowski, F., 2022. Disentangling craving- and valence-related brain responses to smoking cues in individuals with nicotine use disorder. Addict Biol 27, e13083.
Janes, A.C., Farmer, S., Peechatka, A.L., Frederick Bde, B., Lukas, S.E., 2015. Insula-Dorsal Anterior Cingulate Cortex Coupling is Associated with Enhanced Brain Reactivity to Smoking Cues. Neuropsychopharmacology 40, 1561-1568.
Janes, A.C., Frederick, B., Richardt, S., Burbridge, C., Merlo-Pich, E., Renshaw, P.F., Evins, A.E., Fava, M., Kaufman, M.J., 2009. Brain fMRI reactivity to smoking-related images before and during extended smoking abstinence. Exp Clin Psychopharmacol 17, 365-373.
Janes, A.C., Smoller, J.W., David, S.P., Frederick, B.D., Haddad, S., Basu, A., Fava, M., Evins, A.E., Kaufman, M.J., 2012. Association between CHRNA5 genetic variation at rs16969968 and brain reactivity to smoking images in nicotine dependent women. Drug Alcohol Depend 120, 7-13.
Kaag, A.M., Reneman, L., Homberg, J., van den Brink, W., van Wingen, G.A., 2018. Enhanced Amygdala-Striatal Functional Connectivity during the Processing of Cocaine Cues in Male Cocaine Users with a History of Childhood Trauma. Front Psychiatry 9, 70.
Karl, D., Bumb, J.M., Bach, P., Dinter, C., Koopmann, A., Hermann, D., Mann, K., Kiefer, F., Vollstadt-Klein, S., 2021. Nalmefene attenuates neural alcohol cue-reactivity in the ventral striatum and subjective alcohol craving in patients with alcohol use disorder. Psychopharmacology (Berl) 238, 2179-2189.

Karoly, H.C., Schacht, J.P., Meredith, L.R., Jacobus, J., Tapert, S.F., Gray, K.M., Squeglia, L.M., 2019. Investigating a novel fMRI cannabis cue reactivity task in youth. Addict Behav 89, 20-28.

Kim, J.I., Lee, J.D., Hwang, H.J., Ki, S.W., Park, I.H., Park, T.Y., 2020. Altered subcallosal and posterior cingulate cortex-based functional connectivity during smoking cue and mental simulation processing in smokers. Prog Neuropsychopharmacol Biol Psychiatry 97, 109772.

Kleinhans, N.M., Sweigert, J., Blake, M., Douglass, B., Doane, B., Reitz, F., Larimer, M., 2020. FMRI activation to cannabis odor cues is altered in individuals at risk for a cannabis use disorder. Brain Behav 10, e01764.

Kushnir, V., Menon, M., Balducci, X.L., Selby, P., Busto, U., Zawertailo, L., 2013. Enhanced smoking cue salience associated with depression severity in nicotine-dependent individuals: a preliminary fMRI study. Int J Neuropsychopharmacol 16, 997-1008.

Le, T.M., Zhornitsky, S., Zhang, S., Li, C.R., 2020. Pain and reward circuits antagonistically modulate alcohol expectancy to regulate drinking. Transl Psychiatry 10, 220.

Li, X., Chen, L., Ma, R., Wang, H., Wan, L., Bu, J., Hong, W., Lv, W., Yang, Y., Rao, H., Zhang, X., 2020. The neural mechanisms of immediate and follow-up of the treatment effect of hypnosis on smoking craving. Brain Imaging Behav 14, 1487-1497.

Malcolm, R., Myrick, H., Li, X., Henderson, S., Brady, K.T., George, M.S., See, R.E., 2016. Regional brain activity in abstinent methamphetamine dependent males following cue exposure. Journal of drug abuse 2.

McClernon, F.J., Hutchison, K.E., Rose, J.E., Kozink, R.V., 2007. DRD4 VNTR polymorphism is associated with transient fMRI-BOLD responses to smoking cues. Psychopharmacology (Berl) 194, 433-441.

McClernon, F.J., Kozink, R.V., Rose, J.E., 2008. Individual differences in nicotine dependence, withdrawal symptoms, and sex predict transient fMRI-BOLD responses to smoking cues. Neuropsychopharmacology 33, 2148-2157.

Mondino, M., Luck, D., Grot, S., Januel, D., Suaud-Chagny, M.F., Poulet, E., Brunelin, J., 2018. Effects of repeated transcranial direct current stimulation on smoking, craving and brain reactivity to smoking cues. Sci Rep 8, 8724.

Moran-Santa Maria, M.M., Hartwell, K.J., Hanlon, C.A., Canterberry, M., Lematty, T., Owens, M., Brady, K.T., George, M.S., 2015. Right anterior insula connectivity is important for cue-induced craving in nicotine-dependent smokers. Addict Biol 20, 407-414.

Myrick, H., Anton, R.F., Li, X., Henderson, S., Randall, P.K., Voronin, K., 2008. Effect of naltrexone and ondansetron on alcohol cue–induced activation of the ventral striatum in alcohol-dependent people. Archives of general psychiatry 65, 466-475.

Myrick, H., Li, X., Randall, P.K., Henderson, S., Voronin, K., Anton, R.F., 2010. The effect of aripiprazole on cue-induced brain activation and drinking parameters in alcoholics. J Clin Psychopharmacol 30, 365-372.

Park, M.S., Sohn, J.H., Suk, J.A., Kim, S.H., Sohn, S., Sparacio, R., 2007. Brain substrates of craving to alcohol cues in subjects with alcohol use disorder. Alcohol Alcohol 42, 417-422.

Potvin, S., Lungu, O., Lipp, O., Lalonde, P., Zaharieva, V., Stip, E., Melun, J.P., Mendrek, A., 2016. Increased ventro-medial prefrontal activations in schizophrenia smokers during cigarette cravings. Schizophr Res 173, 30-36.

Ray, S., Hanson, C., Hanson, S.J., Bates, M.E., 2010. fMRI BOLD response in high-risk college students (Part 1): during exposure to alcohol, marijuana, polydrug and emotional picture cues. Alcohol Alcohol 45, 437-443.

Regier, P.S., Monge, Z.A., Franklin, T.R., Wetherill, R.R., Teitelman, A., Jagannathan, K., Suh, J.J., Wang, Z., Young, K.A., Gawrysiak, M., Langleben, D.D., Kampman, K.M., O'Brien, C.P., Childress, A.R., 2017. Emotional, physical and sexual abuse are associated with a heightened limbic response to cocaine cues. Addict Biol 22, 1768-1777.

Rubinstein, M.L., Luks, T.L., Moscicki, A.B., Dryden, W., Rait, M.A., Simpson, G.V., 2011. Smoking-related cue-induced brain activation in adolescent light smokers. J Adolesc Health 48, 7-12.

Schulte, M.H.J., Kaag, A.M., Boendermaker, W.J., van den Brink, W., Goudriaan, A.E., Wiers, R.W., 2019. The effect of N-acetylcysteine and working memory training on neural mechanisms of working memory and cue reactivity in regular cocaine users. Psychiatry Res Neuroimaging 287, 56-59.
Shi, Z., Jagannathan, K., Padley, J.H., Wang, A.L., Fairchild, V.P., O'Brien, C.P., Childress, A.R., Langleben, D.D., 2021. The role of withdrawal in mesocorticolimbic drug cue reactivity in opioid use disorder. Addict Biol 26, e12977.
Sjoerds, Z., van den Brink, W., Beekman, A.T., Penninx, B.W., Veltman, D.J., 2014. Cue reactivity is associated with duration and severity of alcohol dependence: an FMRI study. PLoS One 9, e84560.
Versace, F., Engelmann, J.M., Jackson, E.F., Costa, V.D., Robinson, J.D., Lam, C.Y., Minnix, J.A., Brown, V.L., Wetter, D.W., Cinciripini, P.M., 2011. Do brain responses to emotional images and cigarette cues differ? An fMRI study in smokers. Eur J Neurosci 34, 2054-2063.
Vollstadt-Klein, S., Kobiella, A., Buhler, M., Graf, C., Fehr, C., Mann, K., Smolka, M.N., 2011. Severity of dependence modulates smokers' neuronal cue reactivity and cigarette craving elicited by tobacco advertisement. Addict Biol 16, 166-175.
Vollstadt-Klein, S., Wichert, S., Rabinstein, J., Buhler, M., Klein, O., Ende, G., Hermann, D., Mann, K., 2010. Initial, habitual and compulsive alcohol use is characterized by a shift of cue processing from ventral to dorsal striatum. Addiction 105, 1741-1749.
Walter, M., Denier, N., Gerber, H., Schmid, O., Lanz, C., Brenneisen, R., Riecher-Rossler, A., Wiesbeck, G.A., Scheffler, K., Seifritz, E., McGuire, P., Fusar-Poli, P., Borgwardt, S., 2015. Orbitofrontal response to drug-related stimuli after heroin administration. Addict Biol 20, 570-579.
Wang, W., Zhornitsky, S., Zhang, S., Li, C.R., 2021. Noradrenergic correlates of chronic cocaine craving: neuromelanin and functional brain imaging. Neuropsychopharmacology 46, 851-859.
Zhang, S., Zhornitsky, S., Angarita, G.A., Li, C.R., 2020. Hypothalamic response to cocaine cues and cocaine addiction severity. Addict Biol 25, e12682.
Zhornitsky, S., Zhang, S., Ide, J.S., Chao, H.H., Wang, W., Le, T.M., Leeman, R.F., Bi, J., Krystal, J.H., Chiang-shan, R.L., 2019. Alcohol expectancy and cerebral responses

to cue-elicited craving in adult nondependent drinkers. Biological psychiatry: cognitive neuroscience and neuroimaging 4, 493-504.
